# Modeling of Aryl Hydrocarbon Receptor Pathway Intrinsic Immunometabolic Role using Glioblastoma Stem Cells and Patient-Derived Organoids

**DOI:** 10.1101/2022.03.17.484756

**Authors:** Fumihiro Watanabe, Ethan W. Hollingsworth, Yeshavanth Banasavadi-Siddegowda, Jenna M. Bartley, Steven T. Sizemore, Lianbo Yu, Maciej Pietrzak, Paula Schiapparelli, Vinay Puduvalli, Balveen Kaur, Alfredo Quiñones-Hinojosa, Jaime Imitola

**Author notes:** These authors contributed equally to this work (Fumihiro Watanabe, Ethan W. Hollingsworth).

## Abstract

The intrinsic genetic program of glioblastoma (GBM) stem cells is critical for tumor evolution and recurrence. We recently identified intrinsic phenotypes and immune-like genetic programs of GBM organoids (GBMO)^1^ from patient derived glioblastoma stem cells (GSCs), replicating genomic, metabolic, and cellular aspects of GBM *in vivo*. Aryl hydrocarbon receptor (AHR), a ligand-activated transcription factor, is a key regulator of infiltrating immune cells in gliomas^2, 3^ and associated with poor prognosis, but its role in GSC biology is unknown^2^. Here, we show that AHR is a patient-specific regulator of the glioma intrinsic gene program in GSCs and GSC-derived GBMO that are enriched for AHR. We find that AHR is required for GSC self-renewal, GBMO expansion, radial glia-like cell proliferation, and expression of immune mediators seen in the mesenchymal subtype. CRISPR-Cas9 genetic ablation and pharmacological inhibition revealed that AHR regulates genes linked to intrinsic immunity, proliferation, and migration in GBMO. Genomic analysis of GBMO treated with AHR inhibitors identified expression signatures and candidate markers associated with survival of gliomas. Our work defines the glioma intrinsic function of AHR in a model of early GBM formation, offering a rationale for clinical exploration of a potential ‘two-hit’ target of both GBM cells and infiltrating immune cells in patients with GBM expressing high levels of AHR.

## Introduction

The search for vulnerabilities in glioblastoma (GBM) has long been centered on reducing glioblastoma stem cell (GSC) proliferation and more recently on bolstering immune cell infiltration, particularly of T cells. The latter of which normally target the tumor itself before undergoing exhaustion due to tumor-secreted or surface-expressed molecules that inactivate antitumoral T cell responses^4^. One such molecule released by GBM is kynurenine, a ligand for the transcription factor, aryl hydrocarbon receptor (AHR)^5^, in infiltrating immune cells. Activation of AHR in T cells leads in to depression of T cells^2^ and activation of tumor associated macrophages (TAM) with CD39 activation, CD8 dysfunction, and secretion of chemokines^3^. While the pathways that suppress immune cell activation are becoming increasingly defined, the intrinsic transcriptional mechanisms by which GBM cells, including GSCs influence infiltrating immune cell types are still not fully understood^6, 7^. One hypothesis is that GSCs cells have an intrinsic genetic program that regulates cell-to-cell and short-range paracrine interactions with immune cells. Support for this hypothesis stems from the glial origins of GBM, as normal glial and neural progenitors can mount a facultative immune response during neuroinflammation^8, 9^. Indeed, GBM cells can secrete cytokines, and express T cell inhibitors and class I MHC molecules^4^. Thus, determining how the intrinsic properties of GBM contribute to immune cell crosstalk is critical for understanding how these highly lethal tumors evade the immune response and identifying targetable pathways.

One major difficulty in identifying GBM intrinsic programs is separating the contribution of the glial tumor compartment from other components of the tumor microenvironment (TME), including immune cells, pericytes, and endothelial cells. By the time a GBM is clinically detected, the interactions of GBM with surrounding cells of the TME are already established. In turn, the experimental window into understanding the evolution of these GBM-immune cell interactions is lost, thereby preventing their identification. Modeling these interactions *in vitro* has been hampered by the lack of translational models that offer 3D resolution.

Recently, patient-derived tumor organoids – 3D cultures that mimic complex tissue formation – have been established, including from GBM, offering opportunities to address these questions. GBM organoids derived from intact pieces of the tumor, for instance, allow for the study of established interactions in the TME but are less suitable to model the initial interactions due to the simultaneous presence of GBM and multiple types of stromal cells^10^ or to understand the intrinsic immune potential of GBM. By contrast, organoids derivate from GSCs^11^ and without said stromal cells may model the early organization of 3D tumor tissue from GSCs and the intrinsic program as it evolves, before engaging in immune cell crosstalk. Using this patient-derived GBM organoid model from GSCs, heretofore GBMO, we previously defined an intrinsic GBM gene expression program, which despite the absence of classical immune infiltrates, was enriched for immune-like genes derived from tumoral GSC. The critical role of AHR in controlling immune cell activation is well-defined, however the intrinsic role of AHR and its downstream targets in GSCs and GBM biology, is less understood beyond its association with pathways such us TGF-β^12^.

In searching for transcriptional regulators of the GSCs and GBMO intrinsic immunity program, we found patient-specific expression of the aryl hydrocarbon receptor (AHR) pathway. Taking advantage of the GSCs and the GBMO system without stromal-cell-expressed AHR, here we use an integrative approach to delineate the functional role of AHR in a 3D model of the initial stages of GBM in GSCs. We find that the recently discovered AHR-associated gene, interleukin-4-induced-1 (*IL4I1*) an upstream activator of AHR in gliomas implicated in patient survival^13^ and *AHR* are co-expressed in GBMO in a patient-specific manner. We demonstrate AHR controls self-renewal and proliferation of GSCs, and GBMO evolution by transcriptional regulation of genes associated with tumor immunology, metabolism, stemness, and the mesenchymal phenotype. Finally, we expand on the role of kynurenine finding that drives the expression of AHR-regulated genes important for self-renewal and immunity in GSCs and GBMOs. Collectively, our study expands on the role for AHR in early GBM formation, identifying it as a regulator of intrinsic GSCs biology, as modeled by GSC-derived GBMO, and reinforces the future targeting of this pathway to reduce intrinsic GBM stemness and phenotypes associated with GBM progression^2^.

## Results

### Identification of the GBM intrinsic AHR pathway in patient derived GSCs and GBMOs

Recently, we performed a deep phenotypic characterization of GBMO from patient-derived GSCs, and found that they recapitulate GBM *in vivo* with unique self-organization, cancerous metabolic states, and mutational landscapes^11^ (**Figure 1a**). Despite being devoid of immune cells, transcriptomic analysis across GBMO revealed an immune-like molecular program, enriched in genes encoding cytokines, antigen presentation and processing, T-cell receptor inhibitors, and interferon effectors that was particularly enriched in an important population of GBM progenitors^11^. We reasoned that this intrinsic glioma gene expression may be under tight transcriptional control. To identify glioma-intrinsic immune-like upstream regulators, we performed differential expression analysis comparing four different patient- GSCs derived GBMO and NSC-derived organoids and focused on molecules that were shared among the GBMOs (**Fig 1b and Supplementary Fig 1)**. We found 84 converging differentially expressed immune associated genes among GBMOs and then specifically examined the expression of transcription factors (TFs), reasoning that TFs could dictate the GBMO-intrinsic molecular landscape (**Fig. 1b**). This approach, together, with an unbiased upstream TF prediction analysis using all genes upregulated in GBMOs (**Fig. 1c**), led us to identify several heterogeneously expressed candidates, including *ZBTB38*, *SMAD6*, and *NFE2L2.* We then asked what cells express these TFs in the GBM microenvironment *in vivo*. We found that many of these TFs are expressed both by various GBM tumoral cell types and by infiltrating immune cells *in vivo,* using published single cell RNA-seq data (**Fig 1d)**. The top candidate was *AHR* (p<1x10^-65^), which is highly expressed in two of the GBMO and is an immunometabolic regulator that has been studied in the context of infiltrating immune cells in gliomas^2^ but not in GSCs or GBMO. Accordingly, we examined the expression of AHR-associated genes, *IDO1*, *IDO2*, *TDO2,* and *IL4L1*. *IL4L1* encodes for an enzyme recently shown to be required for *AHR* expression and involved in cancers, including GBM^13^. We found that only *IL4L1* was consistently co-expressed with *AHR* in the same GBMO (**Fig 1e**), in line with recent findings^13^. These data support a patient-specific role for the *IL4L1-AHR* pathway in GBMO and provide impetus to further interrogate the functional role of AHR as a regulator of the GBM intrinsic genetic program and GSCs.

**Figure 1.**
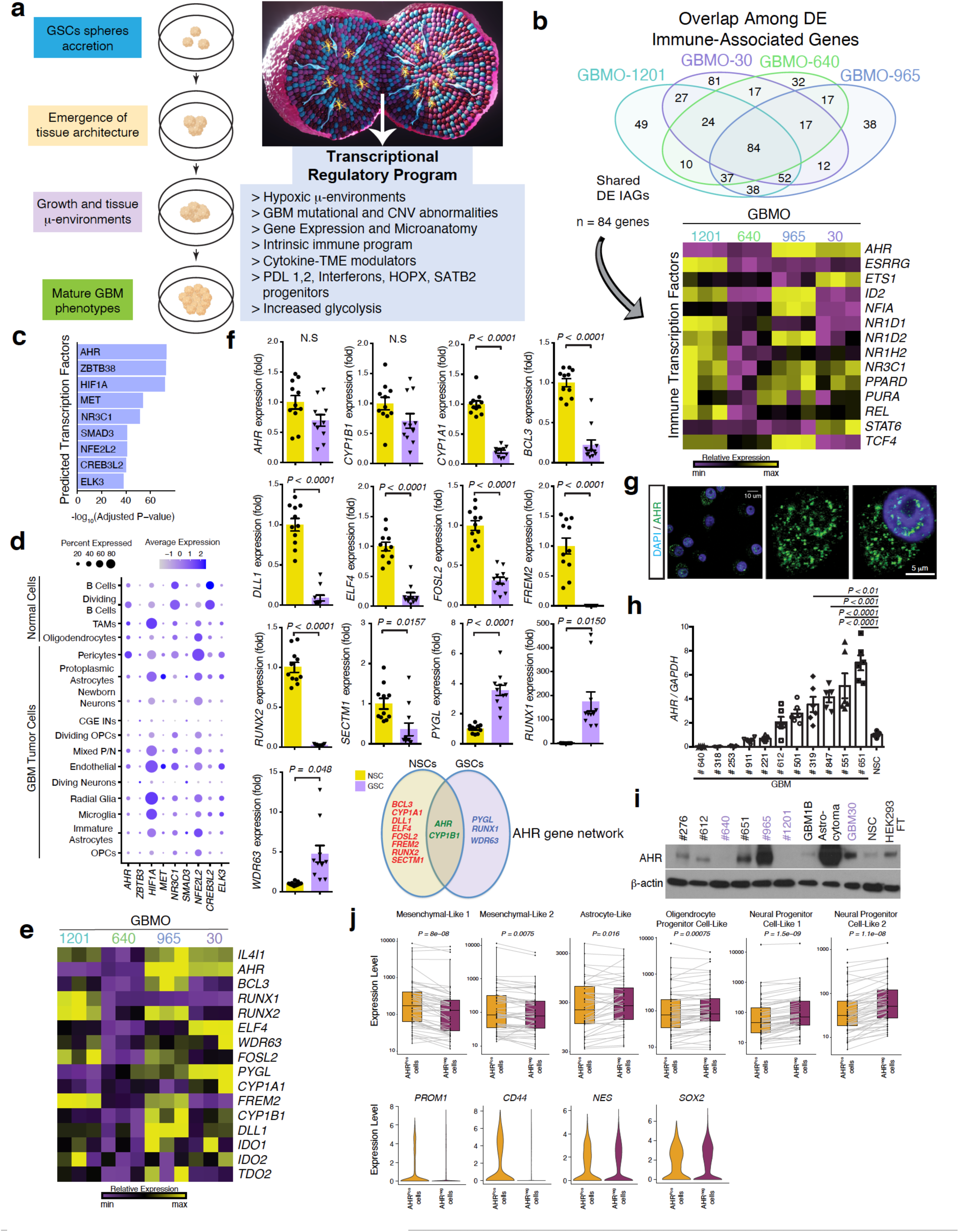
GBMO intrinsic program analysis reveals enrichment of immune genes and patient specific AHR expression. **(a)** Schematic workflow for identifying genes comprising the GBMO intrinsic program during the in vitro evolution. **(b)** Bioinformatic approach for identifying targetable immune-related transcription factors (TFs) with heatmap (right) depicting shared differentially expressed TFs in each GBMO, Briefly, we focused on the shared top 33% of expressed genes from each GBMO line. **(c)** Predicted upstream TFs, as inferred from genes belonging to largest GBMO-enriched gene cluster. **(d)** Single cell RNAseq plot of the expression of TF identified in GBMOs in in vivo GBMs**. (e)** Heatmap depicts the expression of AHR network genes in each GBMO line. **(f)** Confirmatory qPCR of AHR network gene expression in human NSCs (yellow) and GBM30-derived GSCs (purple). Venn diagram indicates differential gene expression in NSCs, compared to GSCs. Two-tailed unpaired Student’s t-test; *n*= 3 independent experiments. Bar graphs are presented as means ± SEM. (*BCL3, P <* 0.0001; *CYP1A1, P<* 0.0001; *DLL1, P < <* 0.0001; *ELF4, P <* 0.0001; *FOSL2, P <* 0.0001; *FREM2, P<* 0.0001; *PYGL, P<* 0.0001; *RUNX1, P=* 0.0150; *RUNX2, P<* 0.0001; *SECTM1, P =* 0.0157; *WDR63, P <* 0.048) **(g)** Distribution of AHR in GSCs. Immunocytochemical images of GBM30 cultures labeled with AHR (green) and DAPI (blue). Scale bars, 10 and 5 μm. **(h)** *AHR* mRNA expression in GSCs procured from several patients with GBM and NSCs. One-way ANOVA with Dunnett’s correction, relative to NSC; n = 3 independent experiments. Data presented as means ± SEM. (GBM319, *P* < 0.01; GBM847, *P* < 0.001; GBM551, *P* < 0.0001; GBM651, *P* < 0.0001). **(i)** Expression of AHR by western blot in GSC lines, human NSCs, and HEK293 FT cells (positive control). **(j)** Pair-wise comparison of the expression of subtype-specific genes between AHR-positive and AHR-negative cell populations. Representative violin plots for top marker genes in Mesenchymal-Like clusters (*CD44* and *PROM1*) and Neural Progenitor Cell-Like clusters (*NES* and *SOX2*) between both cell populations. Wilcoxon signed-rank test. scRNA- seq dataset of patient GBM samples is from Neftel et al^15^

To determine the importance of this pathway in GSCs and GBM biology, we first examined the expression of *AHR* and eleven AHR target genes (procured from published ENCODE ChIP-seq^14^ and microarray datasets^2^) in our GBMO transcriptome data. We found that AHR target genes were expressed and retained patient specificity in AHR-positive GBMO-30 and -965 (**Fig 1e).** Confirmatory qPCR of NSC- and GBM30 GSC-derived tumorspheres revealed an equal expression of canonical AHR genes, and distinct upregulation of AHR target genes in NSCs, with exception of *PYGL*, *RUNX1*, and *WDR63,* which were increased in GSCs (**Fig 1f)**. While these AHR target genes are confirmed to be expressed across multiple tumor compartments of GBM in the Ivy GAP spatial RNA-seq dataset (**Supplementary Fig. 2a, b)** and overexpressed in GBM in The Cancer Genome Atlas (TCGA) **(Supplementary Fig. 3a, b)**, interestingly, only the GSC-upregulated AHR genes, *AHR, RUNX1,* and *WDR63*, are associated with mortality in GBM (**Supplementary Fig. 2c**). Through immunostaining of GSCs, we confirmed that AHR expression in GSCs localized to the nucleus and cytoplasm, supporting its role as a TF (**Fig. 1g**). Finally, to extend our findings to other patient lines, we evaluated *AHR* in multiple GSC lines using qPCR and immunoblots (**Fig. 1h, i, and Supplementary Fig.4**). These data demonstrate that *AHR* expression varies widely among patient GSCs, in what can be characterized as *AHR*-proficient and *AHR-*deficient GSCs and GBMO.

To confirm our GBMO findings in GBM *in vivo*, we made use of a published single-cell RNA-sequencing dataset of resected GBM samples and asked in which cell types AHR is expressed in the tumor microenvironment *in vivo*^15^. Using their designated GBM cell types, we found AHR-positive cells to be most highly enriched for mesenchymal-like gene expression, compared to neural progenitor-, astrocyte-, or oligodendrocyte progenitor-like gene expression (**Fig 1j**). To determine whether AHR is expressed in GSCs *in vivo*, we examined the expression of several GSC marker genes in AHR-positive cells^16^. Indeed, GSC genes like *PROM1* (also known as CD133) and *CD44* were dramatically upregulated in AHR-positive cells while neural stem cell-related markers like *SOX2* and *NES* were downregulated, matching what we observed in GBMO, indicating that AHR is expressed in glioblastoma stem cells in vivo and in vitro GSCs and GBMOs.

### AHR promotes GSC self-renewal *in vitro* and GSC-driven tumor growth *in vivo*

To study the role of AHR in GBMO, we initially focused on AHR function in the GBMO-forming GSCs. We first assessed the effects of pharmacological agents on *AHR* levels using the AHR agonist, 6- formylindolo (3,2-b) carbazole (FICZ), and antagonists CH223191 and 6, 2’, 4’-trimethoxyflavone (TMF). We found no differences in *AHR* expression upon treatment with AHR-targeting drugs (**Fig. 2a**). We treated *AHR*-proficient GBM-30-GSCs cultured as tumorspheres with FICZ to activate *AHR* and evaluate its impact on GSC self-renewal. FICZ increased the number of tumorspheres in a dose-dependent manner (**Fig. 2b,c**). Conversely, treatment with CH223191 decreased tumorsphere formation across a range of sizes (**Fig. 2d-f**). To further support these observations, we performed self-renewal assays, which revealed AHR antagonism by CH223191 decreases GBM-30 self-renewal in GBM-30 in a dose- dependent manner (**Fig. 2g**). We replicated these findings in the mouse GBM cell line GL261N grown as tumorspheres (**Supplementary Fig. 5 a, b**).

**Figure 2.**
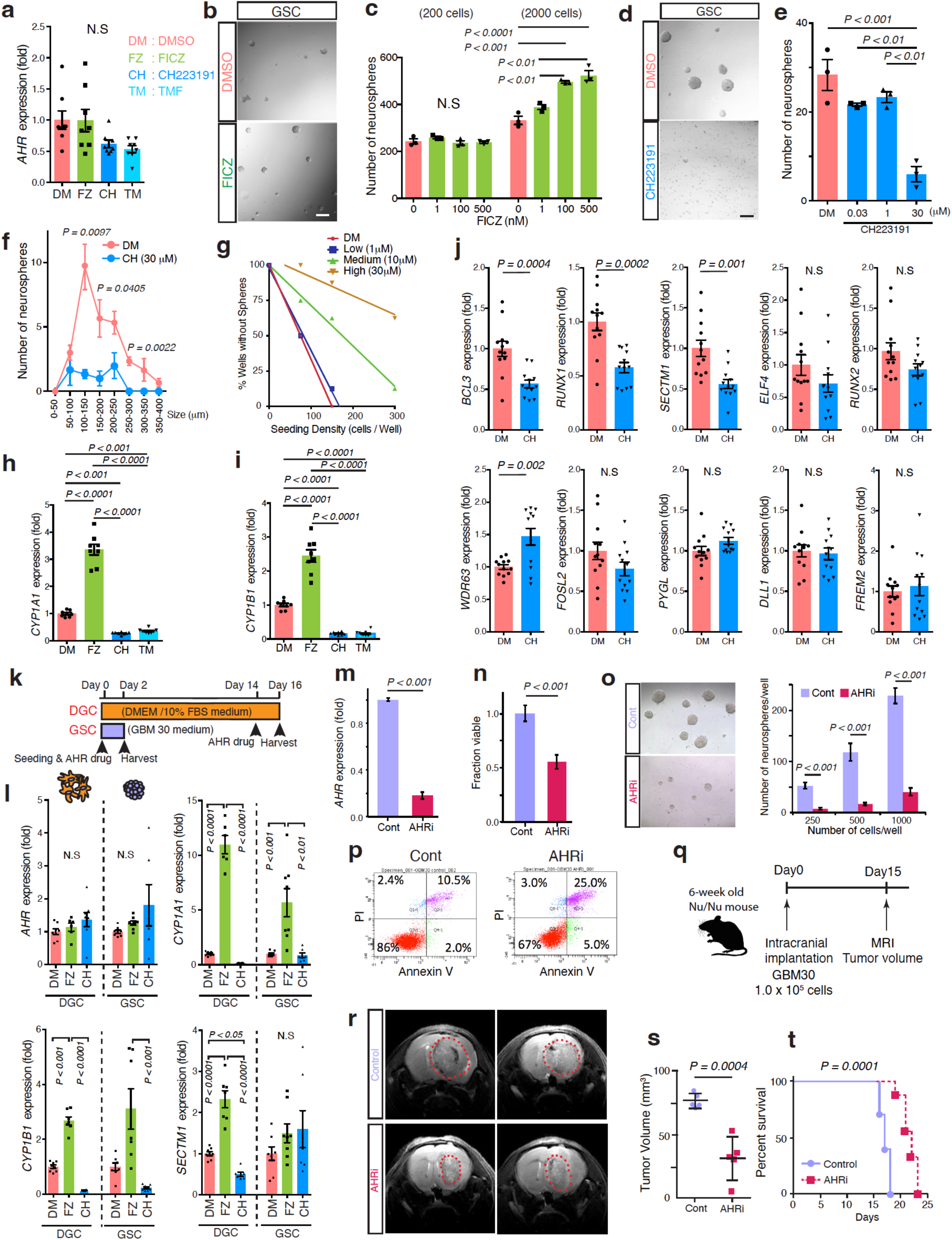
AHR mediates tumorsphere growth *in vitro* and *in vivo*. **(a)** Quantitative RT-PCR for AHR in GBM30 GSCs after treatment with AHR agonist, FICZ (6-formylindolo (3,2-b) carbazole) (FZ), and antagonists, CH223191 and TMF (6,2’,4’-trimethoxyflavone) (TM), or vehicle control DMSO (DM). **(b)** Representative images of GSC tumorsphere formation after DMSO- and FICZ-treatment of GSCs. Scale bar, 100 μm. **(c)** Quantitative analysis of GSC tumorsphere number with increasing FICZ concentration (0, 1, 100, 500 nM) for 7 days when seeded at 200 or 2000 cells/mL. One-way ANOVA with Tukey’s test; *n = 3* independent experiments. Data presented as means ± SEM. (2000 cells: 0 vs. 100 nM, *P* < 0.001; 0 vs. 500 nM, *P* < 0.0001; 1 vs 100 nM, *P* < 0.01; 1 vs. 500 nM, *P* < 0.01). **(d)** Sample images of GSC neurosphere formation after DMSO and AHR antagonist CH223191 (30 μM) treatment for 7 days. Scale bar, 100 μm. **(e)** Quantitative analysis of GSC tumorsphere number with increasing CH223191 concentration (0, 0.03, 1, 30 μM) for 7 days. One-way ANOVA with Tukey’s test; *n = 3* independent experiments. Data presented as mean ± SEM. (DM vs. 30 μM, *P* < 0.001; 0.03 vs 30 μM, *P* < 001; 1 vs. 30 μM, *P* < 0.01). **(f)** Size of GSC tumorspheres when treated with DMSO (red) and CH223191 (30 μM, blue). Two-tailed Student’s t-test; *n* = 3 independent experiments. Data presented as means ± SEM. (100-150 μm, *P =* 0.0097; 150-200 μm, *P* = 0.0405; 250-300 μm, *P =* 0.0022). **(g)** Limited dilution assay depicts clonogenicity of GBM30 GSCs treated with DMSO and increasing CH223191 (1, 10, 30 μM). **(h and i)** Quantitative RT-PCR for *CYP1A1* (h) and *CYP1B1* (i) mRNA after AHR drug treatment in GSCs. One-way ANOVA with Tukey’s test; *n* = 3 independent experiments. Data are presented as mean ± SEM. (*CYP1A1*: DM vs. FZ, *P <* 0.0001; DM vs. CH223191, *P* < 0.001; DM vs. TMF*, P* < 0.001, FZ vs. CH223191, *P <* 0.0001; FZ vs. TMF, *P* < 0.0001. *CYP1B1*: DM vs. FZ, *P <* 0.0001; DM vs. CH223191, *P* < 0.001; DM vs. TMF*, P* < 0.001, FZ vs. CH223191, *P <* 0.0001; FZ vs. TMF, *P* < 0.0001). **(j)** mRNA expression of AHR network genes in DMSO and CH223191-treated GSCs. Two-tailed Student’s t-test; *n* = 3 independent experiments. Data presented as means ± SEM. (*BCL3*, *P* = 0.0004; *RUNX1*, *P =* 0.0002; *SECTM1, P* = 0.001; *WDR63, P* = 0.002). **(k)** Time course, culture conditions, and AHR drug treatment for differentiated glioblastoma cells (DGCs) and non-differentiated GSCs. **(l)** Quantitative RT-PCR for *AHR* and AHR network genes after AHR drug treatment in GSCs and DGCs. One-way ANOVA with Tukey’s test; *n* = 3 independent experiments. Data presented as mean ± SEM. (*CYP1A1* DGC: DMSO vs. FICZ, *P* < 0.0001; FICZ vs. CH223191, *P* < 0.0001. *CYP1A1* GSC: DMSO vs. FICZ, *P* < 0.001; FICZ vs. CH223191, *P* < 0.01. *CYP1B1* DGC: DM vs. FICZ, *P* < 0.001. *CYP1B1* GSC: FICZ vs. CH223191, *P* < 0.001. *SECTM1* DGC: DMSO vs. FICZ, *P* < 0.0001; DMSO vs. CH223191, *P* < 0.05; FICZ vs. CH223191, *P* < 0.0001). **(m)** Expression of *AHR* mRNA in GSCs transfected with AHR siRNA (AHRi), relative to mock-infected GSC tumorspheres (Cont). Two-tailed unpaired Students’ t-test; *n* = 3 independent experiments; *P* < 0.001. **(n)** Fraction of viable AHRi- and Cont-transfected GSC tumorspheres. Two-tailed unpaired Student’s t-test; n =3 independent experiments; *P* < 0.001. **(o)** Representative images (10X) show the size and number of AHRi- and Cont-transfected GSC tumorspheres. Two-tailed unpaired Student’s t-test; *n* = 3 independent experiments. (250 cells/ well, *P* < 0.001; 500 cells/well, *P* < 0.001; 1000 cells/well, *P* < 0.001). **(p)** Apoptotic assays and cell cycle analysis of AHRi- and Cont-transfected GSCs. **(q)** Experimental procedure for measuring changes in tumor size and mouse survival after cell transplantation. **(r)** T2-weighted coronal MRI slices depicting largest tumor section at day 15 post implantation. **(s)** Tumor volumes in mice implanted with Cont (*n* = 3) and AHRi (*n* = 3) GSCs. Two-tailed unpaired Student’st t-test; *P* = 0.0004. **(t)** Kaplan-Meier survival plots of mice implanted with Cont (*n =* 6) and AHRi (*n* = 6) GSCs. Mantel-Cox analysis; *P* = 0.0001.

Next, we asked how pharmacological manipulation of AHR affected its downstream gene expression. qPCR of FICZ-treated GSCs showed an upregulation of canonical AHR genes *CYP1A1* and *CYP1B1*, while both antagonists (CH223191 and TMF) decreased their expression (**Fig. 2h, i**). We expanded our assay to the full host of canonical genes and determined AHR inhibition decreased the expression of several AHR targets, including the anti-apoptotic genes *BCL3* and *RUNX1*, and the immune-associated gene *SECTM1*, but increased *WDR63* expression (**Fig. 2j**).

As evidence suggests the critical TFs orchestrating glioblastoma formation may differ among GSCs and their differentiated progeny, it will be most favorable to identify TFs that are constitutively active during tumor evolution (in both GSCs and differentiated cells), allowing for a double therapeutic “hit” in the process of targeting tumor progression^17^. To assess whether AHR effectors remain functionally expressed in the late stages of GSC differentiation, we differentiated GSCs for 14 days into differentiated GBM cells (DGCs) and compared the expression of canonical genes between GSCs and DGCs. mRNA levels of AHR effector genes, *CYP1A1*, *CYP1B1*, *SECTM1*, *BCL3*, *RUNX2*, *FOSL2*, *ELF4*, and *DLL1*, were predominantly increased in DGCs (**Supplementary Fig. 5c**). We then examined whether pharmacological manipulation of AHR in GSCs and DGCs would yield differential effects on canonical gene expression, as shown in **Figure 2k**. While treatment with FICZ and CH223191 produced heterogeneous changes in several genes (**Supplementary Fig. 5d**), it was highly effective in altering *CYP1A1* and *CYP1B1* mRNA levels in GSCs and their differentiated progeny (**Fig. 2l**). This suggests GBM cells are responsive to AHR-mediated transcriptional changes, irrespective of their differentiation stage, making AHR an ideal candidate to target both GSCs and DGCs.

To confirm these pharmacological observations in an alternative experimental paradigm, we transfected GSCs with small-interfering RNA (siRNA) to target *AHR* transcripts. Successful knockdown of AHR was confirmed in AHR-inhibited GSCs (AHRi) via qPCR (**Fig. 2m**). Functionally, we tested the effect of AHR knockdown on GSC self-renewal and found a decreased number of tumorspheres formed by AHRi GSCs after 7 days of culture compared to mock-transfected GSCs (Cont) (**Fig. 2o**). We determined that reduced self-renewal in AHRi-treated cells was caused, in part, by increased cell death of GSCs (**Fig. 2n**). We further verified this result with Annexin-V staining, which showed that the decreased viability of AHRi GSCs was due to increased apoptosis after 7 days (**Fig. 2p**). To further support this finding, we performed an unbiased screening of protein phosphorylation in AHRi and Cont GSCs. We observed a significant increase in the phosphorylation of GSK3α/β, WNK1, and p53 at three different positions: S46, S392, and to a lesser extent S15 (**Supplementary Fig. 6a**). Similar results were obtained upon pharmacological depletion of AHR activity with CH223191, particularly in the phosphorylation of p53 (S46) and GSK3α/β (**Supplementary Fig. 6b**). These results suggest that AHR inactivation is detrimental to GSCs renewal, which in part, occurs by preventing the phosphorylation of p53.

To determine whether our *in vitro* observations of the role of AHR in GSCs extend to an *in vivo* context, we implanted Cont and AHRi GSCs from GBM30 – a highly malignant mesenchymal GBM cell line with extensive validation *in vivo*^18–20^ – by intracranial injection into mice (**Fig. 2q**). GSCs were cultured for 48 hours after siRNA treatment before being injected into mice with a similar number of live cells for each group. MRI scans of tumor-bearing mice 15 days post implantation revealed striking differences in tumor volume. AHRi-implanted mice had noticeably smaller tumors than their Cont- implanted counterparts (**Fig. 2r, s**). Additionally, AHR knockdown conferred a significant survival advantage, as the median survival of AHRi-implanted mice was 22 days, opposed to 17 days in the control group (**Fig. 2t**). These findings underscore the relevance of our *in vitro* findings, supporting a direct link between AHR and GSC self-renewal *in vivo*.

### AHR regulates in vitro evolution and intrinsic properties of GSC-derived GBMO

Next, we examined the functional role of AHR in the 3D tumor-like environment provided by GSCs forming GBMO by evaluating the effect of AHR activation and inactivation on GBMO phenotypes from the formation until 4 weeks later. We evaluated histological organization and gene expression (**Fig 3a**). First, we cultured AHR-proficient GBMO-30 with FICZ, CH223191, or DMSO (vehicle control) for 4 weeks. We found that compared to DMSO, GBMO cultured with FICZ were larger with multiple nodules, whereas CH-treated GBMO were significantly smaller (**Fig 3b**). These differences were supported by histopathology of GBMO. FICZ-treated GBMO showed nuclear atypia and an expanded ∼500 μm active border of proliferation (**Fig. 3c**). CH223191-treated GBMO, by contrast, exhibited a marked decrease in cellularity with areas devoid of cells in a background of extracellular matrix (**Fig. 3b, c**). To further confirm these findings, we stained and quantified the number of proliferating Ki-67^+^ and apoptotic A-Cas^+^ cells in each GBMO condition. Quantification revealed increased proliferation in FICZ-treated and increased apoptosis and decreased cell proliferation in CH223191-treated organoids, relative to DMSO cultures (**Fig. 3d, e**), indicating that targeting AHR limits GBMO growth in part by reducing proliferation and increasing apoptosis.

**Figure 3.**
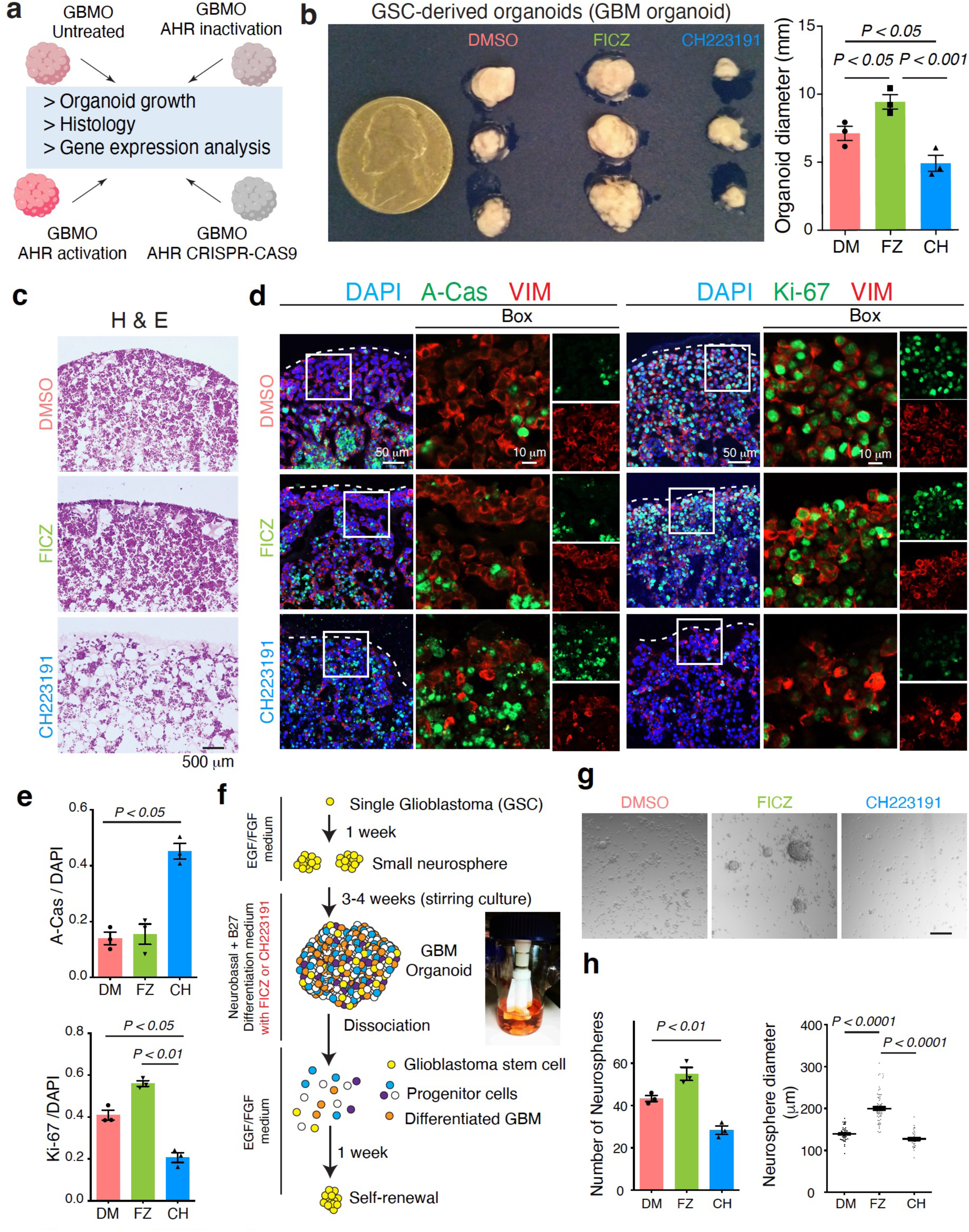
Bidirectional impact of AHR targeting in GBMO. **(a)** Schematic diagram for studying patient824 derived glioblastoma in the GBM organoid model using methods of AHR activation by agonist, inactivation by antagonist, and gene knockout by CRISPR-Cas9. **(b)** Representative images and quantification of GBMO-30 size after treatment with AHR agonist/antagonist after 4 weeks in stirring culture. One-way ANOVA with Tukey’s test; *n* = 3 independent experiments. Data are presented as mean ± SEM. (DMSO vs. FICZ, *P* < 0.05; DMSO vs. CH223191, *P* < 0.05; FICZ vs. CH223191, *P* < 0.001.) **(c)** Hematoxylin & Eosin (H&E) staining in GBMO-30 after treatment with DM or exogenous AHR ligands. Scale bar, 500 μm. **(d)** Effect of AHR manipulation on GBMO proliferation (Ki-67) and apoptosis (A-Cas). Sample confocal images of A-Cas (left) and Ki-67 (right) in GBMOs treated with FICZ or CH223191 compared to control (DMSO). Note the increased A-Cas and decreased Ki-67 staining in CH223191- compared to DMSO- and FICZ-treated GBMO. Scale bars, 50 and 10 μm. **(e)** Quantitative analysis for A-Cas^+^ and Ki-67^+^ cells in AHR drug-treated GBMO. One-way ANOVA with Tukey’s test; *n* = 3 independent experiments. Data presented as mean ± SEM. (A-Cas: DMSO vs. CH223191, *P* < 0.05. Ki-67: DMSO vs. CH223191, *P* < 0.05; FICZ vs. CH223191, *P* < 0.01.) **(f)** Scheme of self-renewal analysis of GBMOs at 3 weeks after AHR drug treatment. **(g)** Representative images of GSC tumorsphere formation after 7 days using cells isolated from 3-week-old GBMOs after DMSO and AHR drug treatment. Scale bar, 100 μm. **(h)** Quantification of tumorsphere number and diameter, indicating self-renewal capacity. One-way ANOVA with Tukey’s test; *n* = 3 independent experiments. Data presented as mean ± SEM. (Number: DMSO vs. CH223191, *P* < 0.01. Diameter: DMSO vs. FICZ, *P* < 0.0001; FICZ vs. CH223191, *P* < 0.0001.)

In the complete tumor ecosystem, GSCs act as a quiescent, self-renewing population of cells resistant to standard chemotherapies^21, 22^. To rule out that the observed differences in organoid size were from preferential targeting only DGCs, we performed a tumorsphere formation assay from GBMO, that we have used for determining neural stem cell proliferation from *in vivo* tissue^23, 24^ (**Fig. 3f)**. After three weeks of treatment with FICZ or CH223191, we dissociated GBMOs and cultured the same number of viable cells in FGF-2 and EGF media to test their ability to form tumorspheres (**Fig. 3g**). Notably, quantitative analysis for tumorsphere number and diameter indicated self-renewal capacity increased in FICZ- and decreased in CH223191-treated organoids (**Fig. 3h**). We confirmed these observations at the molecular level, observing reduced expression of both progenitor (PAX6) and differentiated cell (TBR1, SATB2) markers in CH223191-treated GBMO (**Supplementary Fig. 7a,b**). Together, these results suggest that inhibiting AHR simultaneously targets both GSCs and DGCs in GBMO.

To determine whether AHR plays a ubiquitous role in regulating GBM biology, we expanded our growth analysis to GBMOs derived from additional patient GSCs with varying AHR expression. We first performed a limiting dilution assay to determine self-renewal capacity in each GSC line and found that AHR-proficient (AHR^Pos^) GSCs from GSC30 and GSC965 show more self-renewal activity than AHR- deficient (AHR^Neg^) GSCs from GSC1201 and GSC640 (**Fig. 4a**). To interrogate the role of AHR in self-renewal, we cultured GSCs from AHR^Pos^ GSC965 and AHR^Neg^ GSC1201 as neurospheres in the presence of DMSO or CH223191. CH223191 treatment of AHR^Pos^ GSCs led to decreased neurosphere formation and size, while AHR^Neg^ GSCs showed no different after CH223191 (**Fig. 4b, c**). To extend our observations in an alternative assay, we examined the effects of AHR antagonism on the limited dilution assay of AHR^Pos^ and AHR^Neg^ GSCs at different concentrations, we also tested its effect on human NSCs.

**Figure 4.**
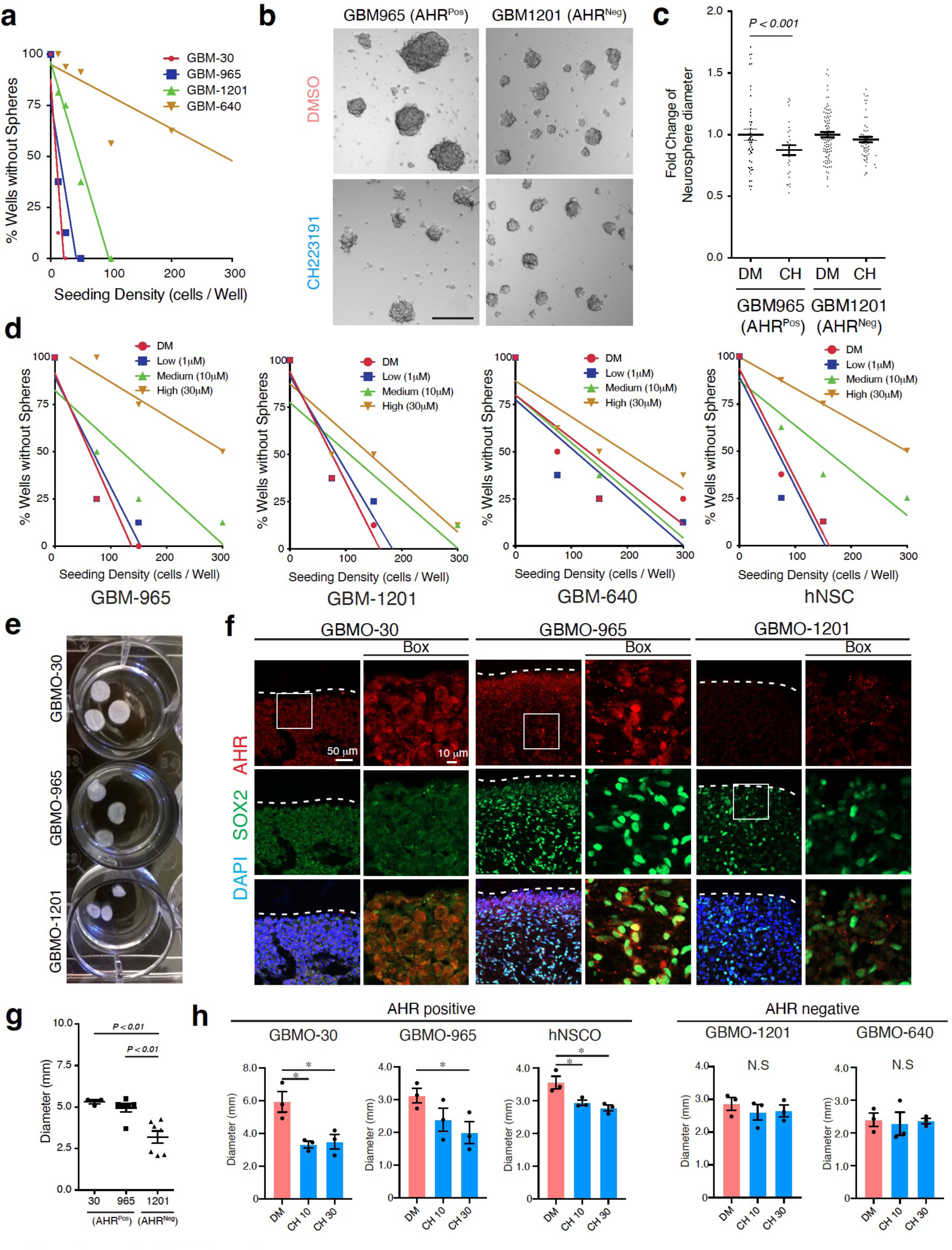
Endogenous AHR levels associate with responses to targeting AHR in GBMO. **(a)** Limited dilution assays depicting clonogenicity of all patient-derived GSCs. (**b**) Bright-field images of GSC tumorsphere formation for GSC965 (AHR^Pos^) and GSC1201 (AHR^Neg^) after CH223191 treatment (30 μM). Scale bar, 100 μm. **(c)** Fold-change plots of AHR^Pos^ and AHR^Neg^ GSCs tumorsphere diameter after DMSO and CH223191 treatment. Two-tailed unpaired Student’s t-test; *n* = 3 independent experiments. Data presented as mean ± SEM. (GSC965: DMSO vs. CH223191, *P* < 0.001). **(d)** Comparison of clonogenicity by limited dilution assays of each patient derived GSC with increasing CH223191 concentration (0, 0.03, 1, 30 μM). **(e, f**) Representative images and quantification of each patient derived GBMO diameter. One-way ANOVA with Tukey’s test; *n* = 3 independent experiments. Data are presented as mean ± SEM. (30 vs. 1201, *P* < 0.01; 965 vs. 1201, *P* < 0.01.). (**g**) Sample images of SOX2 and AHR immunostaining in each GBMO-30, -965, and -1201. Scale bars, 10 and 50 μm. (**h**) Quantification of diameter for each GBMOs (GBMO-30, -965, -1201, -640) and hNSCO after 4 weeks in stirring culture with CH223191 treatment. One-way ANOVA with Tukey’s test; *n* = 3 independent experiments. Data are presented as mean ± SEM.

We observed a significant reduction in self-renewal in AHR^Pos^ lines GSC-30 (**Fig 2g**), GSC-965 and NSCs (**Fig 4d**). Next, we generated GBMO from these AHR^Pos^ and AHR^Neg^ GSCs. Despite initiating each organoid with a similar number of GSCs, AHR^Neg^ GBMO were visibly smaller after 4 weeks, compared to AHR^pos^ as measured by microscopy and confocal images (**Fig. 4e-g**). We then generated GBMO from each GSC line, as well as NSC-derived organoids, and targeted them with AHR using CH. Like in GSCs, only AHR^Pos^ organoids showed differences in GBMO size in response to CH (**Fig 4h**). Altogether, these results suggest that while AHR indeed drives GBMO growth, it only does so in AHR-proficient cells, which in turn renders them susceptible to AHR targeting in a dose-dependent manner and expand our observations to additional GBMO organoids.

### Inhibition of AHR pathway in GBMO targets self-renewal, immune, and radial glial programs

Having demonstrated that AHR is a functional regulator of GBMO-965 and -30, we next focused on the molecular targets of AHR in GBMOs. To do this, we performed a genome-wide transcriptome analysis in AHR-proficient GBMO-30 treated with DMSO-, FICZ-, and CH223191 because of its highly malignant profile in animals and mesenchymal phenotype mimicking a highly malignant GBM. Differential expression analysis (FC > ±2.0, *p* < 0.05) of FICZ-treated versus DMSO control identified 59 DEGs and showed an upregulation of known AHR effector genes, including *CYP1A1* and *CYP1B1*, as well as several other immune-related oncogenes, such as *JUN*, *MSTIL*, and *SS18*^25^ (**Fig. 5a, b and Supplementary Table 1**). In contrast, treatment with CH223191 robustly altered the GBMO-30 transcriptome, resulting in 771 DEGs, including several prognostically important chemokines and interleukins in GBM, such as *CCL20*^32^ and *IL1B.* Network analysis demonstrated the central role of AHR in DEGs (**Fig. 5a, b and Supplementary Table 2**). More notably, CH223191-treatment decreased the expression of genes associated with neural stemness, such as *HES1*^26^, and GBM stemness and the mesenchymal subtype, like *ALDH1A3* and *CD44*^27, 28^ (**Fig. 5b,d**). We performed confirmatory qPCR of several key DEGs and GO and pathway analysis determined that FICZ and CH223191 treatments led to differential expression of genes involved in tumor cell migration and proliferation in GBM-30 (**Supplementary Fig. 8a**). We extended these findings to GBMO-965 using GBM30 as control in *SERPINEB2* and *TIPARP*, two key AHR target genes (**Supplementary Fig. 8 b,c)**.

**Figure 5.**
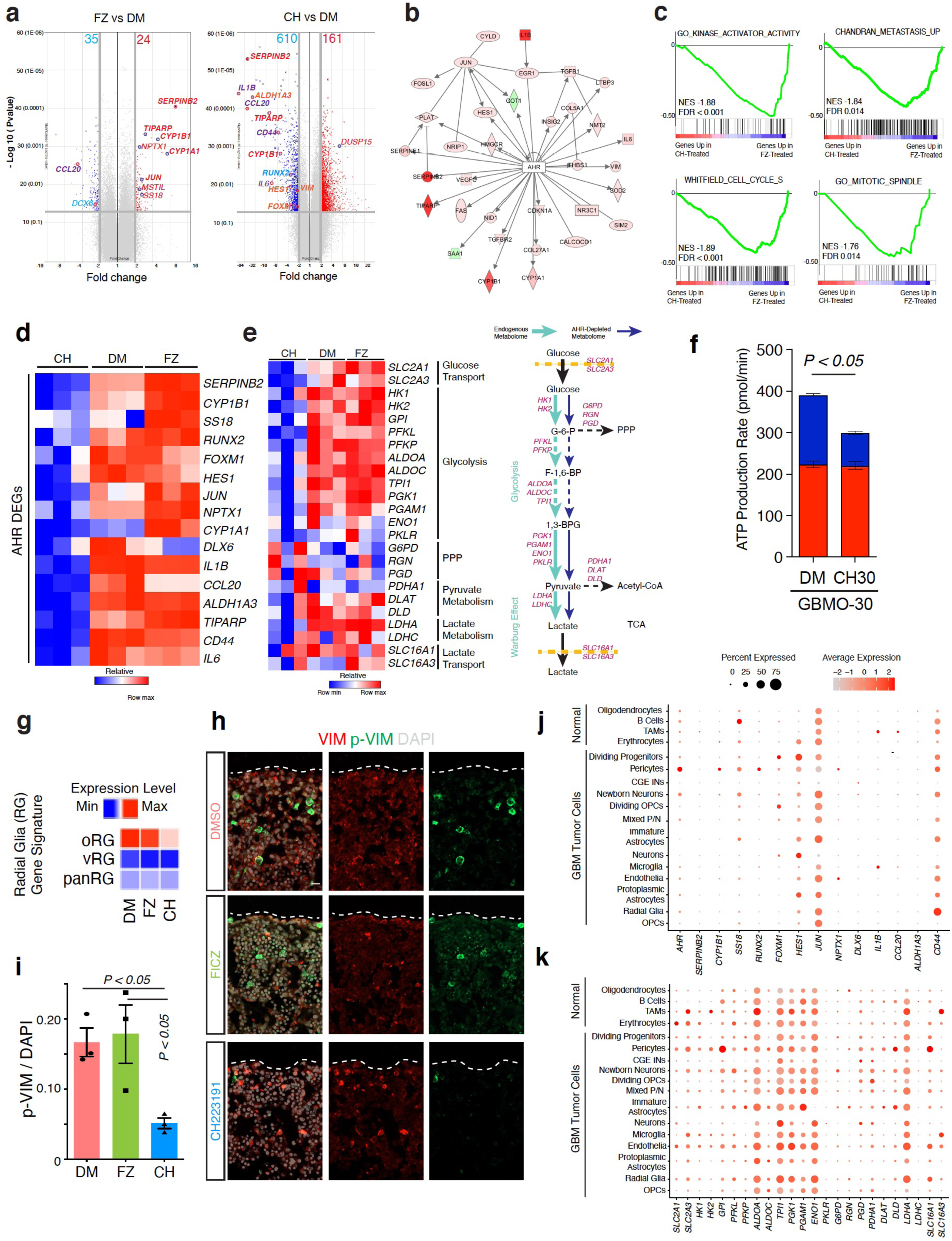
AHR activation and inhibition in GBMO modulates stemness, radial glial and immunometabolic genetic programs. **(a)** Volcano plots of gene expression between DMSO- vs. FICZ- treated (left) and DMSO- vs. CH223191-treated (right) GBMOs. Fold-change (log_2_), x-axis; -log_10_(adj. *P- value*), y-axis. **(b)** Inferred biological network predicts AHR to be upstream of the observed transcriptional changes. Molecules are colored by differential expression (fold change). Red, upregulated; green, downregulated. **(c)** Representative GSEA enrichment plots show cell cycle and cancer-related ontologies enriched in FICZ-treated GBM organoids. False Discovery Rate (FDR) and normalized expression score (NES) calculated by GSEA software. **(d)** Heatmap of differentially expressed genes (DEGs) related to stemness and immunity in DMSO- (*n* = 3), FICZ- (*n* = 3), and CH223191-treated (*n* = 3) GBMOs. **(e)** Metabolism gene profiles of CH223191-, DMSO-, and FICZ-treated GBMO-30. Schematic summary of biochemical pathways indicates that depleting AHR attenuates the Warburg Effect and restores PPP flux. Arrow thickness represents relative expression. 1, 3-BPG, 1, 3-Bisphoshoglycerate; F- 1,6-BP, Fructose-1, 6-Bisphosphate, G-6-P, Glucose-6-Phosphate; PPP, Pentose Phosphate Pathway; TCA, Tricarboxylic Acid. (**f**) Seahorse XF Real-Time ATP Rate Assay to calculate mitochondrial ATP (Mito ATP) and glycolytic ATP (Glyco ATP) production rates in CH223191-treated GBMO-30 and controls. Total ATP: p= 0.026, Glyco ATP: p=0.7707, Mito ATP: p<0.0001. **(g)** Heatmap showing average expression level in GBMO-30 for outer radial glia (oRG), ventricular radial glia (vRG) and pan- RG gene signatures after AHR drug treatment. **(h)**Representative images of immunostaining for Vimentin and the mitotic radial glia marker, p-Vimentin, in AHR drug-treated GBMO-30 samples. Scale bar, 20 μm. **(i)** Quantification of p-Vimentin^+^outer radial glia-like cells in GBMO-30 after treatment with AHR drugs. One-way ANOVA with Tukey’s test. Data are presented as mean ± SEM. (DMSO vs. FICZ, *P* < 0.05; FICZ vs. CH223191, *P* < 0.05.) **(j)** In vivo expression of DEG in single GBM cells from scRNAseq database from Muller et al^54^. (**k**) In vivo expression of metabolic genes in single GBM cells from scRNAseq database from Muller et al^54^

On a global scale, using Gene Set Enrichment Analysis, we determined that transcripts upregulated by FICZ were enriched for oncogenic processes, such as metastasis, mitosis, and kinase activity (**Fig. 5c**). Next, we investigated how AHR inactivation impacted GBMO-30 metabolic gene expression. Importantly, inhibiting AHR with CH223191 treatment disrupted the Warburg Effect and shifted GBMO toward a non-malignant metabolic state by diminishing glycolytic and lactate metabolism while concomitantly restoring the pentose phosphate pathway; this was confirmed by Seahorse functional analysis of GBMO-30 treated with CH223191 and DMSO (**Fig. 5e, f and Supplementary Fig. 9 a-d**).

In our prior characterization of GBMO, we found outer-radial glia-like cells (oRG) with highly migratory morphologies^11^. As such, we asked how AHR inhibition impacted the oRG gene signature found in human GBM^29^. AHR antagonism significantly decreased the expression of oRG genes in GBMO, while preserving the gene signatures of ventricular radial glia (vRG) and panRG (**Fig. 5g**). To examine these observations at the cellular level, we stained treated GBMO-30 for p-VIM a marker for mitotic radial glia. Strikingly, CH223191 treatment led to a marked decrease in the number of p-VIM^+^ oRG-like tumor cells (**Fig. 5h, i**). We next examined functional pathways altered by AHR activation and inactivation showing consistent differential regulation of AHR, TGFβ, cancer, inflammation and migration molecular pathways (**Supplementary Fig. 10 a,b**). We confirmed immunological associated genes involved in cell migration. Using qPCR, we determined that *CCL20*, which varied significantly by patient tumor, was significantly upregulated in GBM30 GSCs, while *CCR6*, the receptor for CCL20, was decreased (**Supplementary Fig. 10c,d**). At the protein level, we found that AHR antagonism decreased CCL20 levels, as measured by ELISA in two independent GSC lines treated with AHR inhibitors (**Supplementary Fig. 10e**).

To determine the cell types that would be targeted by AHR antagonism in GBM *in vivo*, we turned a single-cell RNA-seq dataset with normal and pathogenic cell type annotation (ref). Using their cell type annotations, we examined the expression of AHR network and CH223191-downregulated genes. Many of these genes, including *CD44,* were highly expressed in radial glia-like and astrocytic tumor cells, as well as some infiltrating immune cells, like B cells (**Fig 5j**). Likewise, metabolic genes are highly expressed in the glial compartment of GBM *in vivo* (**Fig 5k**). These findings provide evidence for the AHR and metabolic signaling pathway being active in radial glia-like and astrocytic tumor cells *in vivo*.

Next, to investigate how the transcriptomic changes of GBMO induced by AHR inhibition could affect signatures of GBM *in* vivo, we examined the expression of DEGs using TCGA and Ivy GAP^30^. Interestingly, many of the identified DEGs correlated with AHR at the transcriptional level in GBM *in vivo* (**Supplementary Fig. 11**). We then correlated the identified DEGs with tumor microanatomy. FICZ- activated genes are predominantly expressed in the perinecrotic compartment of GBM, that is modeled by hypoxic areas in the GBMO as previously shown^11^ (**Supplementary Fig. 12 a,b**). CH223191-inhibited genes, on the other hand, were not only highly expressed in the perinecrotic zone but also at the leading edge and microvasculature which are associated with hypoxic microenvironments, that in vivo are populated by vascular and migratory tumoral cells (**Supplementary Fig. 12 c-e and Table 3**).

To confirm the role of AHR as a transcriptional regulator of GBMO suggested by the gene expression dynamics after both activation and inactivation of AHR, we envisioned that AHR was directly binding at target gene promoters. Indeed, by using ENCODE ChIP-seq data for AHR, we observed robust binding upstream to several of these genes (**Fig. 6a**). To functionally show that these genes are targets of AHR, we made use of CRISPR-Cas9 and edited exon 1 of AHR, which encodes for its DNA- binding domain, in GBM-30 GSCs (**Fig. 6b**). We confirmed successful DNA editing using Sanger sequencing and that AHR protein levels were ablated in by Western blot analysis (**Fig. 6c**, **d** and **Supplemental Fig 13**). We then performed a limited dilution assay of tumorspheres using WT and AHR CRISPR-edited GBM-30 GSCs and found decreased proliferation in AHR CRISPR-edited GSCs and decreased GBMO size **(Fig. 6 e,f)**. qPCR revealed a decrease in *SERPINEB2*, *TIPARP*, *CYP1B1*, and *ALDH1A3* in AHR CRISPR-edited GSCs, replicating our whole transcriptomic results (**Fig. 6g**). more notably, to further determine a direct, AHR-dependent role in the expression of these genes, we measured their gene expression upon stimulation by AHR agonist in AHR CRISPR-edited cells compared to controls. Upon treatment with FICZ, only control, but not AHR CRISPR-edited GSCs, showed upregulation of genes critical for cancer and stemness, including *SERPINB2*, *TIPARP*, *CCL20*, and *ALDH1A3* (**Fig. 6h**). The same result occurred with addition of the endogenous AHR activator, kynurenine, wherein control, but not AHR CRISPR-edited cells, showed gene upregulation (**Fig. 6i**). Finally, we noted a similar reduction as pharmacological targeting of AHR in the metabolic activation of GBMO from AHR CRISPR-edited GSCs (**Fig. 6j, k** and **Supplemental Fig 9e**). These results indicate that AHR transcriptionally regulates the expression of genes in AHR-proficient GBMO that may be clinically relevant for highly malignant mesenchymal GBM or recurrent GBM with mesenchymal transition, which are SOX2^-^, CD44^+^, and require *ALDH1A3*^31^.

**Figure 6.**
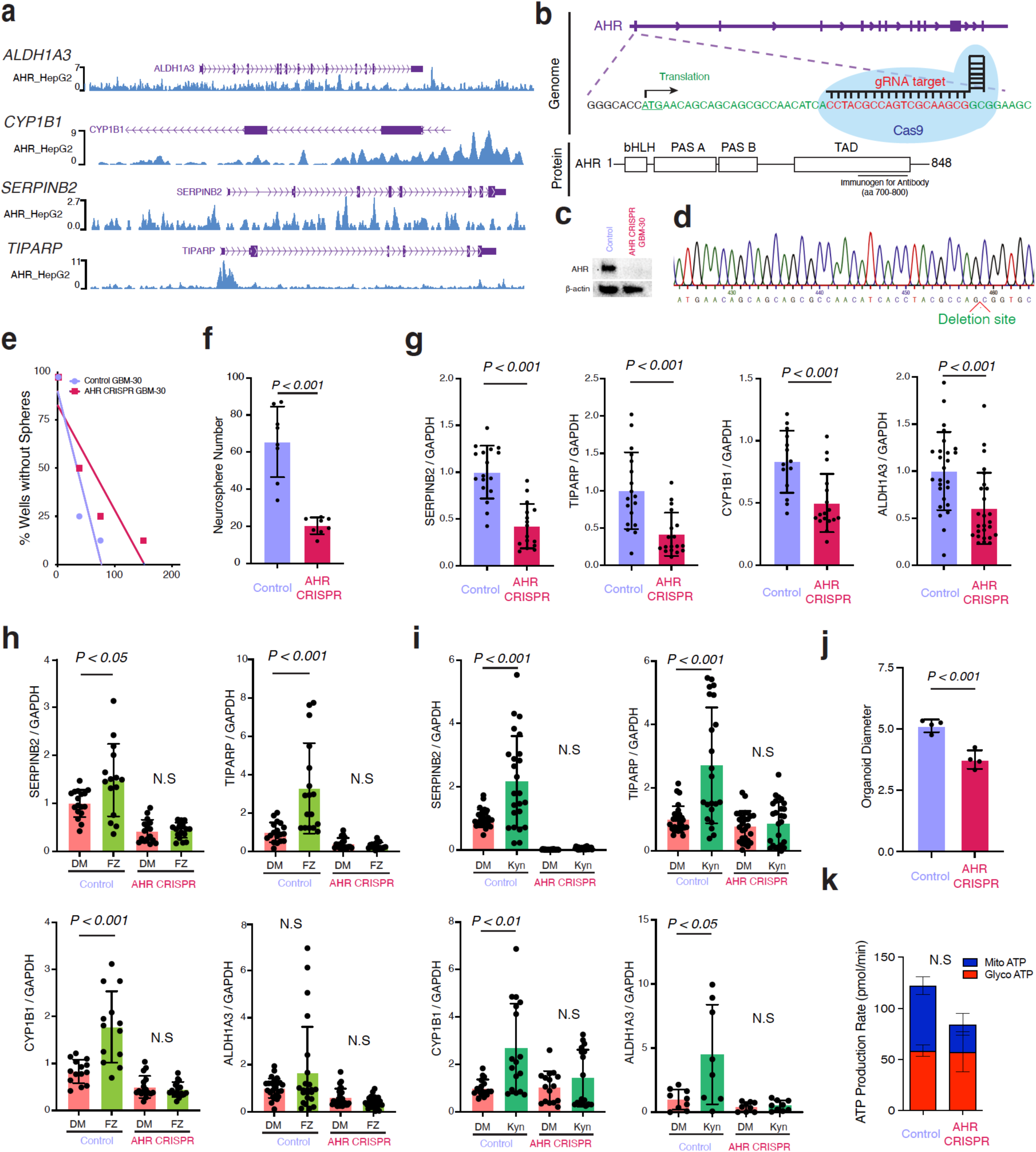
Functional genetic inactivation with CRISPR-Cas9 confirms a role for AHR in the regulation of GBMO target genes. **(a)** WashU Epigenome Browser tracks for AHR genomic occupancy upstream of AHR network genes (*ALDH1A3*, *CYP1B1*, *SERPINB2* and *TIPARP*) using ChIP-seq data in HepG2 cells. **(b)** Genomic and protein location of designed gRNA for CRISPR-mediated genetic targeting AHR exon 1. **(c)** western blot (right) for AHR protein levels in control and CRISPR-treated GBM-30 GSCs. **(d)** Sequencing for target region (left) in CRISPR-treated GBM-30 GSCs. **(e, f)** Limiting dilution (left) and tumorsphere (right) assay for control and CRISPR-GBM-30 GSCs. **(g)** qPCR for mRNA expression of AHR network genes, *SERPINB2*, *TIPARP*, *CYP1B1*, and *ALDH1A3,* in control and CRISPR-treated GBM-30 GSCs. *n =* 3 independent experiments. **(h, i)** Expression of AHR network genes in control and CRISPR-treated GBM-30 GSCs after treatment with FICZ (h) and kynurenine (Kyn, i). **(j)** Size of organoids derived from WT and CRISPR-GBM-30 GSCs. **(k)** Seahorse XF Real-Time ATP Rate Assay to calculate Mito ATP and Glyco ATP production rates from AHR-CRISPR GBMO-30 vs controls: p=0.3669, Mito ATP: p=0.0552, Glycol ATP: p=0.9520.

### An AHR-negative expression signature in GBMOs induced by AHR blockade is associated with prognosis and potential GBM targets

Having previously shown that GBMO preserve transcriptomic alterations seen in GBM *in vivo*, we leveraged our GBMO platform to identify potential GBM molecules of interest based on AHR manipulation. First, we expanded our analysis of the *in vivo* GBM expression level to expression- associated mortality for genes upregulated in GBMO-30 by a 2.5-fold change relative to NSCO. Using this integrated approach, we found 41 genes similarly overexpressed in organoids that matched the *in vivo* GBM that were associated with poor prognosis in either GBM or its subtypes (**Fig. 7a**). Of the 41 identified genes, 19 had not been previously described in GBM or implicated in conferring a significant survival advantage. Importantly, we determined that ten of the 41 genes were downregulated following AHR inhibition in GBMO-30 and significantly associated with survival supporting a role of inactivation of AHR on potential genes that drive mortality in GBM. These include *PLP2*, a gene involved in autophagy and ER stress in GBM^32^; *SGMS2,* which is associated with aggressive breast cancer^33^; *MT1H,* which is a tumor suppressor in liver cancer^34^; and *SERPINE1*, which is associated with several cancers^35^ (**Fig. 7b- e**). These results indicate that GBMO may represent a tool for identifying future molecular targets not only relevant for a specific tumor, but also for GBM in general.

**Figure 7.**
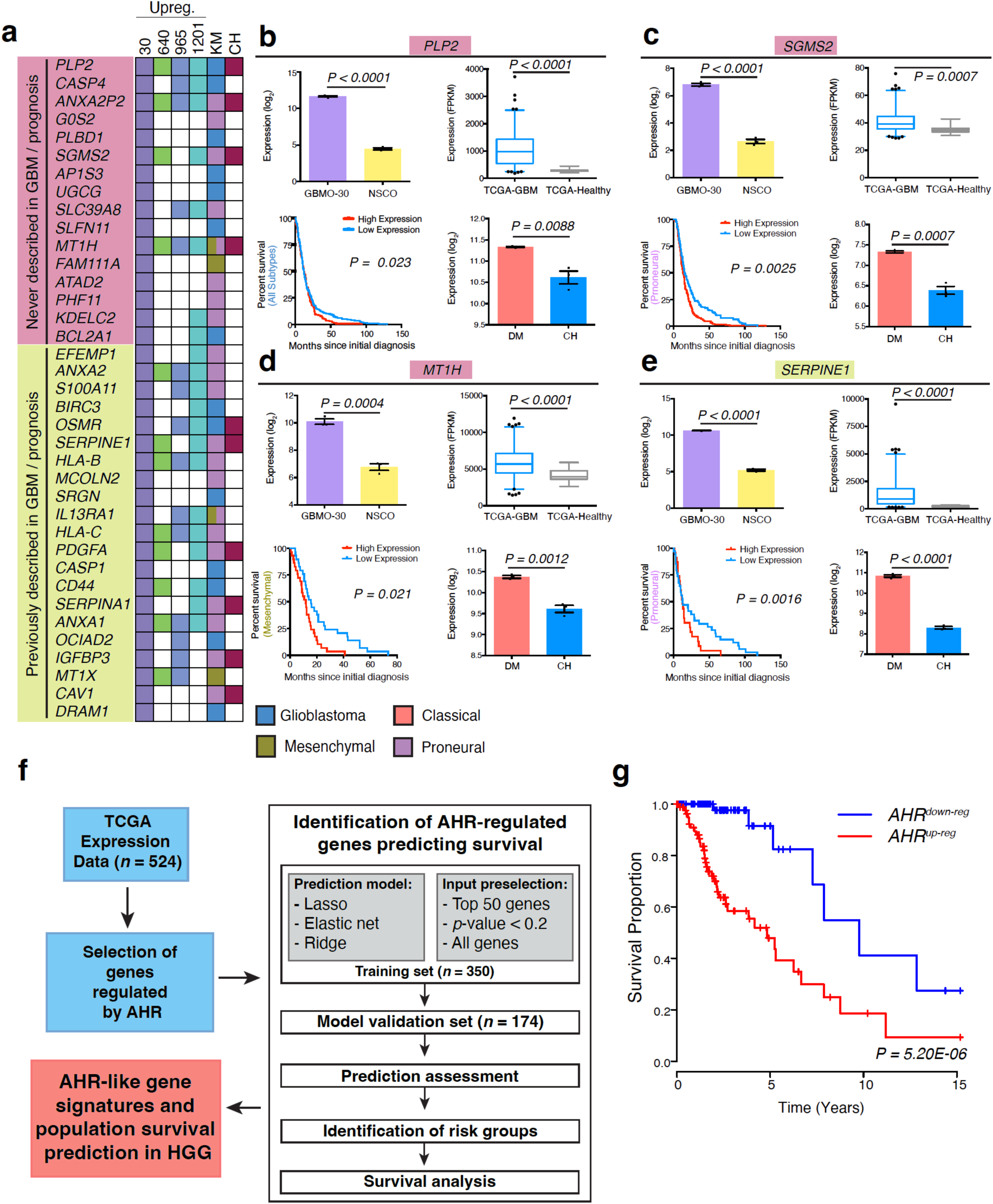
AHR inhibition associates with survival advantage for patients with gliomas. **(a)** Table summarizing the results of the gene expression (OE, overexpression), outcome prediction (KM, Kaplan- Meier), and differential expression by CH223191 treatment (FC<-2) analyses for genes from the upregulated GBMO-30 signature, using publicly available TCGA cohorts. **(b)** Expression of*PLP2* in GBMO-30 compared to NSCO (*P* < 0.0001), in GBM samples versus healthy tissue in the TCGA-GBM database (*P* < 0.0001), and between DMSO- and CH223191-treated GBMO-30 (*P* = 0.0088). Kaplan- Meier plot (*P* = 0.023) also indicates low *PLP2* expression also confers a significant survival advantage all GBM. **(c)** Expression of *SGMS2* in GBMO-30 compared to NSCO (*P* < 0.0001), in GBM samples versus normal tissue in the TCGA-GBM database (*P* = 0.0007), and between DMSO- and CH223191- treated GBMO-30 (*P* = 0.0007). Kaplan-Meier plot (*P* = 0.0025) also indicates low *SGMS2* expression confers a significant survival advantage for the proneural subtype. **(d)** Expression of *MT1H* in GBMO-30 compared to NSCO (*P* = 0.0004), in GBM samples versus normal tissue in the TCGA-GBM database (*P* < 0.0001), and between DMSO- and CH223191-treated GBMO-30 (*P* = 0.00012). Kaplan-Meier plot (*P* = 0.021) also indicates low *MT1H* expression confers a significant survival advantage for the mesenchymal subtype. **(e)** Expression of *SERPINE1* in GBMO-30 compared to NSCO (*P* < 0.0001), in GBM samples versus normal tissue in the TCGA-GBM database (*P* < 0.0001), and between DMSO- and CH223191-treated GBMO-30 (*P* < 0.0001). Kaplan-Meier plot (*P* = 0.0016) also indicates low *SERPINE1* expression confers a significant survival advantage for the proneural subtype. **(f)** Schematic procedure for computational analysis of population survival by AHR expression signature. **(g)** (Top) Survival probabilities of patients with GBM characterized as AHR^up-reg^(red) or AHR^down-reg^ (blue). (Bottom) Expression data is derived from TCGA cohorts. Mantel-Cox test; *P* = 5.20E- 06.

To further assay the value of the GBMO in modeling the effects of inactivating the AHR pathway in highly malignant AHR-competent tumors at a population level, we used mathematical modelling to investigate whether the GBMO expression signature induced by AHR antagonism (AHR^down-reg^) in our AHR positive organoids are relevant for GBM prognosis. For this, we selected a subset of the most downregulated genes detected in the CH-treated GBMO-30 transcriptome and randomly assigned the samples to a training (350 subjects) and testing set (174 subjects) using TCGA data. We then performed training using Cox regression with three models: LASSO, ElasticNet, and Ridge regression. These results were then validated in the testing set and the prediction performance was assessed as satisfactory by using Uno’s estimator of cumulative/dynamic AUC for right-censored time-to-event data (**Fig. 7f**). Collectively, these patient population data indicate that inducing a GBMO AHR^down-reg^ signature in human GBM with AHR^up-reg^ may confer a significant survival advantage to patients (**Fig. 7g**).

## Discussion

Using our previously characterized model of patient derived GBMO^1^, we uncovered that the AHR pathway functions at the intersection of regulating intrinsic immunity, self-renewal, and stemness in GBM. We highlight the integral role of AHR in mediating cell-autonomous GBM self-renewal, metabolism, and immune signaling in GSCs, DGCs, and throughout GBMO evolution. Expansion of our functional experiments to other patient-derived GBMO clarified that AHR is not a ubiquitous driver of GBMO growth, but varies by patient and is associated with the mesenchymal subtype. Identification of AHR status after resection may offer potential opportunities to investigate inactivation of these pathways in treatment-naïve patients. Further extrapolation at the population level showed that inducing an AHR^Neg^ gene signature in an otherwise AHR-proficient GBMO confers a survival advantage. These data suggest that the identification of AHR-positive GBM may support selective clinical trials with AHR inhibitors to induce a favorable AHR signature. As such, future preclinical and clinical trials employing a targeted approach where AHR inhibitors would be delivered locally by convection enhanced delivery^36, 37^ to modulate the growth of GBM may be worth pursuing and will require the interrogation of tumor AHR activity from surgical biopsy and even functional validation with an organoid assay. Thus, in addition to classical pathological and molecular definition, clinical interrogation of GBM could thus benefit from determination of AHR status. Beyond identifying a transcriptional regulator of intrinsic GBM biology, we demonstrate proof-of concept that GBMO can be leveraged to identify a therapeutically targetable, molecular vulnerability of highly malignant GBM. Applying our approach of creating an organoid “avatar” of the patient tumor will help to identify new targets to put forth for potential clinical trials to personalize treatments for GBM and improve their dismal prognosis^4^ (**Supplementary Fig. 14**).

Mounting evidence has shown a clear pro-tumorigenic role for tryptophan metabolites, like kynurenine, in the TME that activate AHR to recruit TAMs and induce anergy in infiltrating cytotoxic T cells^3^. Intriguingly, we also found the AHR pathway to be a critically important to tumor growth, but in a GBM cell-autonomous fashion. Our work thus highlights that AHR plays a dual tumorigenic role, both within the tumor compartment itself and in the TME. It is tempting to speculate that this dual effect in AHR that leads to explosive tumor growth may underlie why these patients have such an aggressive disease course and may explain why an AHR-negative gene signature confers a survival advantage. At the same time, our observation that not all patients with GBM express AHR raises questions into whether such patient-specificity likewise exists for infiltrating cells of the TME. Future studies using co-cultures of patient-derived GBMO and immune cells will allow us to further dissect the immune circuitry of the TME and answer these questions.

GBM are known to hijack endogenous pathways. For instance, GBM cells uptake activity- dependent neuroligins secreted by neurons to bolster their own growth through PI3K-mTOR signaling^38^. This raises two questions in the context of AHR. First, are there other sources of tryptophan metabolites that GBM repurpose to fuel their growth through AHR activation? Prior evidence demonstrates astrocytes and microglia express the necessary enzymes to breakdown tryptophan and secrete kynurenine^39, 40^. Secondly, if GBM hijack the AHR pathway for self-renewal and stemness, it is conceivable that this pathway is already important in the cells of origin of GBM, neural progenitors. Indeed, we found many of the AHR network genes to be highly expressed in NSCs (**Fig. 1f**), and that AHR antagonism decreased their self-renewal and size (**Fig. 2f, g**). Thus, what is the normal role for AHR in neural progenitors and during the tumoral transition? Addressing this question will be important for not only unraveling the molecular drivers of self-renewal in NSCs, but also as we rationally design therapies for GBM that may have potentially deleterious off-target effects on the NSC niche. Our in vitro results showed that AHR inactivation can target normal human NSC stemness, but whether it preferentially targets the hippocampal or periventricular NSC niche in humans is unknown. Future studies will clarify these questions, as it well known that chemo- and radiation therapy can target normal neural progenitors in cancer patients. Beyond NSCs, our work also points to an unrecognized role of the AHR pathway in a novel subpopulation of oRG-like glioblastoma stem cells, which may contribute to the mesenchymal phenotype that we found to be specific for AHR-positive GBMO^41^. These observations demonstrate that patient-derived GSCs have an encoded ability to give rise to highly malignant progenitors in 3D cultures and suggest that GBM growth and invasiveness may rely on malignant hijacking of the oRG-like gene program with mesenchymal features associated with poor prognosis. Future studies should also detail how AHR controls oRG-like populations at a mechanistic level in GBMO.

Given the absence of immune infiltrates and stroma in GBMO cultures, this system may represent an approach for studying the separate influence of GBM on the intrinsic tumor program. We capitalized on this opportunity to determine novel cell-autonomous mechanisms by which AHR contributes to glioma growth, such as regulating cytokine and chemokine gene expression (**Supplementary Fig. 14, 15**) that expand the role of AHR in cancer, including GBM^2, 3, 5, 13^. The potential benefit of these data is underscored by the current disparity in our understanding of tumor immunity between gliomas and other solid tumors; this is evidenced clinically by the success of checkpoint inhibitors in treating many solid tumors^42^, but not the majority of GBM^43^. GBM organoid systems will required continued optimization, including integration of other cell types like vasculature. In this work, we chose a reductionist and robust approach with high efficiency (**Supplementary Fig. 1**) to culture GBM organoids for a period of 30 days, reasoning this a sensible time frame to find a candidate target using a patient-derived GBM organoids to test a potential personalized model, in a disease with 18-month average survival^44^.

In conclusion, our work expands on the role of AHR in GSCs and GBM and supports the notion of several pro-tumorigenic “AHR cellular programs” in GBM. Them being, 1) AHR in infiltrating immune cells affects their ability to target glioma cells; and 2) AHR in glioma cells is responsible for tumor growth, stemness, and interaction with the TME. Our findings uncover the role of AHR in maintaining intrinsic GBM growth through self-renewal, metabolism, and migration, and propose a patient-derived organoid platform that can be used to find patient- and tumor specific-vulnerabilities (**Supplementary Fig. 14**).

## Online Methods

### Culture of glioblastoma stem cells, neural stem cells and generation of organoids

Human GBM stem cells (GSCs) were procured from patients and allocated for human research purposes, per the protocols approved by the Institutional Review Boards (IRB) at The Ohio State University Wexner Medical Center and Mayo Clinic^18, 19^. Confirmatory biological properties as GSC for these lines has been demonstrated elsewhere (Table 1). Patient-derived GSCs were isolated and cultured in Neurobasal, 2% B27 supplement without Vitamin A, Glutamax, Antibiotic-Antimycotic (Thermo Fisher Scientific, Waltham, MA, USA), 50 ng/ml human epidermal growth factor (EGF), 50 ng/ml basic fibroblast growth factor (FGF) (R&D Systems, Minneapolis, MN, USA), and heparin (Millipore Sigma, Billerica, MA, USA) in low-attachment cell culture flasks, as previously described^14^. To induce differentiation, dissociated single GSCs were incubated in differentiation medium including DMEM, Glutamax, 10% FBS, and Anti-Anti (Thermo Fisher Scientific) for 14 days. Human neural stem cells (NSC, H9 hESC-Derived, Invitrogen) were cultured on Geltrex in KnockOut DMEM/F12, 2% StemPro Neural supplement, Glutamax, Antibiotic-Antimycotic (Thermo Fisher Scientific), 50 ng/ml human EGF, 50 ng/ml human FGF in low-attachment cell culture flasks, several subclones were generated for the study. For GBM and NSC organoid formation, dissociated single cells were cultured with GBM or NSC growth media for 4-7 days. GBM or normal neurospheres were transferred to Matrigel droplets (BD Bioscience, San Jose, CA, USA) by pipetting into cold Matrigel on a sheet of Parafilm with 3 mm dimples. These droplets were allowed to gel at 37°C and were subsequently removed from the Parafilm. After 4 days of stationary growth, the tissue droplets were transferred to a spinning bioreactor containing organoid differentiation media including 48% Neurobasal, 48% DMEM/F12, 1% B27 supplement, 0.5% N2 supplement, 1% Glutamax, 0.5% MEM-NEAA, and 1% Antibiotic-Antimycotic^45^. Human iPSCs and iPSC-derived organoids were generated as described previously^45^. All cell lines were handled in accordance with the IBC biosafety practices and relevant ethical guidelines of The Ohio State University College of Medicine, Nationwide Children’s Hospital, and University of Connecticut Health that regulate the use of human cells for research.

### Immunoblot of GSCs

Cells were lysed in 50 mM Tris-HCl (pH 7.4), 150 mM NaCl, 1% NP-40, 0.25% Na-deoxycholate, 5 mM EDTA, and 1 mM PMSF with protease inhibitor cocktail (Thermo Fisher Scientific). Protein samples were separated by sodium dodecyl sulfate–polyacrylamide gels and electroblotted onto polyvinylidene difluoride membrane (GE Healthcare UK, Buckinghamshire, UK). The blots were immunoreacted with rabbit anti-AHR (ab190797, Abcam, Cambridge, MA, USA) and mouse anti-β-actin (sc-47778, Santa Cruz Biotechnology, Dallas, Texas, USA) antibodies. Immunoreacted protein bands were visualized by chemiluminescence (ECL Plus, GE Healthcare UK). Signals were normalized with those of β-actin as an internal standard.

### Histology and immunofluorescence of GBM Organoids

Tissues were fixed in 4% paraformaldehyde for 20 min at 4°C followed by washing in PBS three times for 10 min. Tissues were allowed to sink in 30% sucrose overnight, embedded in OCT compound (Tissue-Tek, Sakura Finetek USA, Torrance, CA, USA), and then cryosectioned at 20 μm. Tissue sections were stained with hematoxylin and eosin, and images were taken with a light microscope (BX41, Olympus, Tokyo, Japan) equipped with a digital camera (DP71, Olympus). For immunofluorescence, sections were incubated successively with 0.25% Triton X-100 and 4% normal horse serum in PBS for 30 min, primary antibodies overnight, and Alexa Fluor 488-, 594-, or 647- conjugated species-specific secondary antibodies for 2 h (Thermo Fisher Scientific). Vectashield Mounting Medium with DAPI (Vector Laboratories, Burlingame, CA, USA) or DAPI (Thermo Fisher Scientific) were used for counterstaining. For single labeling of tissue, the following primary antibodies against the following molecules (immunized species) were used: activated-caspase-3 (559565, BD Pharmingen, San Jose, CA, USA), AHR (ab190797 and ab2769, Abcam), BNIP3 (sc-56167, Santa Cruz, mouse-monoclonal), CD4 (550278, BD Pharmingen), CD68 (556059, BD Pharmingen), CD8 (sc-18913, Santa Cruz), CTIP2 (ab18465, Abcam), HIF1a (ab2185, Abcam), HOPX (HPA030180, Millipore Sigma), Ki-67 (ab16667, Abcam), PAX6 (AB_528427, DSHB, Iowa city, IA, USA), phosphorylated-vimentin (D076-3, MBL, Nagoya, Japan), REELIN (MAB5366, Millipore Sigma), SATB2 (ab92446, Abcam), SOX2 (AF2018, R&D systems), TBR1 (ab31940, Abcam), TBR2 (ab23345, Abcam), and vimentin (sc7557, Santa Cruz). Images were taken with a confocal laser-scanning microscope (LSM800, Carl Zeiss Microscopy GmbH, Jena, Germany).

### Quantitative analysis of immunostaining of GBM organoids

Markers for proliferation and apoptosis (Ki-67 and activated-caspase-3) were quantified and normalized with respect to nuclear DAPI staining, on the outer and inner surface areas of organoid images. Using ImageJ software, quantification of marker staining was performed in 3 equally sized rectangular areas that were overlayed on each outer and inner surface. A minimum of three sections were quantified for each organoid line. Marker staining for mitotic radial glia using p-VIM was quantified in the DMSO-, CH-, and FICZ-treated GBMO-30 organoids, as described above. The length of relative layer thickness was measured using ImageJ software.

### Real-Time Quantitative PCR of GBM organoids

RNAs were extracted from cell cultures with QIAzol reagent and miRNeasy Mini Kit (QIAGEN, GmbH, Hilden, Germany), following the manufacturer’s protocol. cDNAs were obtained from 500 ng of mRNA using the retrotranscription kit (Thermo Fisher Scientific). Quantitative real-time PCR was performed on 1/20 of the retrotranscription reaction using SYBR Green PCR Master Mix (Thermo Fisher Scientific). Primers were designed to amplify 50- to 200-bp fragments; All primer sequences can be found in **Supplementary Table 1.** The qPCR data were assessed using delta-delta CT for evaluating results in the sigmoid region of the amplification curve. For each analysis, samples were normalized by comparison with the housekeeping gene GAPDH. All samples, including “no template” controls, were assayed in triplicate. Each experiment was performed three times with comparable results. Data are expressed as mean ± SEM.

### siRNA cell transfection and assays of cell viability, cell cycle, and apoptosis of GSCs

Control and AHR-specific siRNAs were purchased from GE Dharmacon (Lafayette, CO, USA). siRNAs were transfected using RNAi Max lipofectamine (Thermo Fisher Scientific) into GBM30 cells by the manufacturer’s protocol. Viability of cells was measured using trypan blue dye exclusion assay (Thermo Fisher Scientific). For cell cycle analysis, post-treatment cells were washed with PBS, fixed with ethanol (80%), and stained with 50 μg/mL propidium iodide (MilliporeSigma). Stained cells were analyzed in a BD LSRII (Becton Dickinson, Franklin Lakes, NJ, USA). To measure apoptosis, cells were subjected to the apoptosis assay using an Annexin V-Fluorescein Isothiocyanate (Annexin V-FITC), PI apoptosis/necrosis detection kit (Abcam), and a FITC-conjugated active caspase inhibitor (ApoStat Apoptosis Detection Kit, R&D Systems), as described previously^18^. Briefly, GSC cells were transfected with Control or AHRi. 72 hr later, cells were harvested and incubated with a FITC-conjugated annexin V gl and PI for 45 min at 37°C or incubated with apostat for 30 min, then harvested and read using a C6 flow cytometer. Annexin V/PI was analyzed on FL1-H versus FL2-H scatter plots and active caspases were detected on FL1-H.

### *In vitro* tumorsphere assay after drug treatment

To determine the effect of AHR treatment on neurosphere size of GSCs, cells were cultured at 200-5000 cells/well in 96-well plates, and then cultured in the presence of the AHR agonist 6- Formylindolo -3,2-b- carbazole (FICZ, Abcam), AHR antagonist 6, 2’, 4’-trimethoxyflavone (TMF, MilliporeSigma), and AHR antagonist CH223191 (Millipore Sigma) at each concentration for a week. The number of neurospheres was counted under inverse microscope (U-TBI90, Olympus, Tokyo, Japan). Each experiment was performed three times with comparable results.

### Antibody array for GSCs

Using human phospho-kinase array (R&D), antibody array was conducted on GBM cells. GSCs were transfected with control or AHR siRNA or treated with AHR antagonist CH223191. 72 hr post- transfection, cells were pelleted and lysed with lysis buffer. Membrane sets pre-coated with antibodies were incubated with lysates overnight at 4°C. After overnight incubation, membranes were processed per the previously reported protocol^18, 19^. Relative expression of the proteins was measured with respect to reference spots of the membrane.

### ELISA analysis of GSCs

The inhibitory effect of AHR antagonist CH223191 on CCL20 expression in GSCs was determined with the ELISA assay. GSCs (3 × 10^5^ /well in 24-well plate) were first treated with CH223191 (30 μM) for 24 h. The expression levels of CCL20 in the supernatants were then examined using ELISA kits (Human CCL20 and IL-β ELISA Kit) according to the manufacturer’s instructions (R&D Systems).

### Intracranial implantation of GSCs in mice

6-week old athymic Nu/Nu mice were intracranially implanted with 100,000 GBM30 tumor cells as previously described^18, 19^ that were transfected with control or AHR siRNA. Tumor cells were implanted 2 mm lateral and 1 mm rostral to bregma at a depth of 3 mm. Mice were anesthetized with ketamine/xylazine solution intraperitoneally and were stereotactically stabilized using Kopf murine stereotactics. Scalps were cleaned and parted, and a 1 mm wide injection hole was drilled using a 1 mm burr bit. Tumor cells were implanted intracranially in a volume of 2 μL Hank’s buffered salt solution over 5 minutes with kd Scientific auto-injectors at a rate of 0.4 L/min. Mice were implanted with GBM30 pre- treated with either si-Scrambled (Cont) or si-AHR (AHRi). Groups are labeled GSC-Cont or -AHRi.

### MRI and tumor size measurement

12 female mice (3 per group) were randomly selected for MRI imaging on day 15 post-tumor implantation. T2-weighted MRI images were captured without gadolinium enhancement, and 1 mm width brain slices were analyzed for tumor volume calculations^18^.

### Transcriptome analysis and differential expression of GBM organoids

The GeneChip Human Transcriptome Array 1.0 (also known as Clariom D assays; Affymetrix, Thermo Fisher Scientific Inc.) was used to provide a detailed analysis of the organoid transcriptome. Briefly, 100 ng of total RNA from each of the three samples originally assigned for microarray analysis were used to generate amplified and biotinylated sense-strand cDNA from the entire expressed genome according to the GeneChip WT PLUS Reagent Kit User Manual (P/N 703174, Affymetrix Inc., Santa Clara, CA). cDNA was hybridized to GeneChip Human Transcriptome Array 1.0 for 16 hr in a 45°C incubator, rotated at 60 rpm. After hybridization, the microarrays were washed, and then stained using the Fluidics Station 450 followed by scanning with the Affymetrix GeneChip Scanner 3000 7G, according to manufacturer’s instructions. Raw intensity data was normalized using the quintile normalization of robust multiarray average (RMA) method (performed at the individual probe level). Probes with low- variance were filtered out using the R package genefilter^46^ and annotated to the human genome using the Human Clariom D platform. Transcripts were identified as differentially expressed using the limma package^47^, with a threshold of FDR-adjusted p-value <0.05 and fold change greater than ±2.

### Principal component analysis, hierarchal clustering, and functional annotation of GBMO gene expression

We reduced the dimensionality of the data by performing principal component analysis (PCA) on the organoid microarray datasets using the prcomp function in R (center = TRUE, scale = FALSE), including only the filtered genes with moderate to high variance. To identify potential transcriptional modules based on the co-expression of genes in the organoid dataset, unsupervised hierarchical clustering of differentially expressed genes (DEGs) was performed using average linkage and uncentered Pearson correlation on variably expressed genes, as determined by the varFilter() function in limma. Log2-scaled expression values were centered on the median before performing hierarchical clustering. Heatmaps of clustered differential gene expression were then generated. Differentially expressed gene data and the resulting clusters (FC±2, *p*<0.05) were exported for further functional analysis. We made use of Enrichr, which utilizes the Fisher exact test with multiple hypothesis testing correction^48^, to determine gene ontologies, dysregulated pathways, and predicted upstream transcription factors.

For the AHR-treatment microarray dataset, pathway analysis was also conducted using Enrichr. Core analyses between pairwise group comparisons in the AHR dataset were conducted in Ingenuity Pathway Analysis Software to generate direct molecule-molecule interaction networks. Further, functional annotations, as determined by the core analyses, were downloaded and filtered to contain only the top-most enriched cellular and molecular categories, as defined by IPA. We made use of the treemap package (https://cran.r-project.org/web/packages/treemap/index.html) to generate treemaps of these annotations, which were sized by their *p-*value and include the genes of the most dysregulated network. Bar graph figures were generated using GraphPad Prism software. For gene ontology analysis considering the entire transcriptome data, raw expression data for all transcripts were imported into Gene Set Enrichment Analysis software to generate enrichment scores for the CH-treated and FICZ-treated gene sets in Hallmark, C2.all, and C5.all MSigDB datasets. Exact parameters are as follows: Ontologies were considered significant when FDR > 0.05 and abs (Normalized Enrichment Scores (NES)) > 1.5. R Programming Language was used to perform all other analyses and generate figures^49^.

### Unbiased search strategy for GSC molecular vulnerabilities

To establish an unbiased GBMO-intrinsic genetic program, normalized intensity values for each GBMO line were first filtered to include only the top one-third of highly expressed genes. The resulting gene lists were compared among GBMO lines, and the genes shared by all four lines were the only ones further considered. Gene lists were inputted into Enrichr for enrichment analyses and CIBERSORT^50^ was utilized for *in silico* enumeration of interacting immune cell subsets from GBMO transcriptomic data using default parameters. To further ascertain the immune expression states of each GBMO line, we concentrated on the expression of immune-associated genes, as obtained from ImmPort (http://www.immport.org/immport-open/public/home/home) and InnateDB (http://www.innatedb.com)^51^, using the original DE analyses to ensure all IAGs were encompassed. Plots were produced using ggplot2 package in R. Again, the convergence of the DE IAGs was utilized and then filtered to focus only on known human transcription factors, as specified by http://fiserlab.org/tf2dna_db//index.html. For heatmap generation, data were imported into the online matrix software, Morpheus (https://software.broadinstitute.org/morpheus).

### Subtype and gene signature analysis

The average relative expression of each set of GBM subtype predictor genes (as defined by TCGA^52^ was quantified from the log2-transformed expression values to determine a relative subtype score for defining each GBMO line. Subtype average expression was normalized by the average log2 expression value of all subtype genes.

To examine gene signatures specifically enriched for brain pericytes and vascular endothelia, we utilized a single-cell RNA-seq dataset from mouse brain vasculature and identified the top 500 enriched genes for pericytes and vasculature (capillary endothelial cell (EC), venous EC, and arterial EC), respectively^53^ (http://betsholtzlab.org/VascularSingleCells/database.html). We extracted the expression values of each respective group from GBMO and NSCO to produce heatmaps. Clusters were determined by discrete upregulation patterns by organoid line and ontologies from Enrichr. To determine whether pericyte and vasculature-enriched genes were preferentially included in each Ivy GAP microvasculature cluster, we examined the overlap of these genes with each GBMO microvasculature cluster, using cellular tumor genes as a theoretical null expectation control. Fisher’s least significant difference test followed by the Bonferroni multiple-hypothesis correction was used to determine statistical significance. Finally, we evaluated how manipulation of AHR expression alters the relative expression of the different radial glia (RG) signatures described by Pollen *et al*., 2015^29^: ventricular RG (vRG), outer RG (oRG), and panRG. Relative expression of oRG, vRG, and panRG genes within the AHR transcriptome dataset was quantified and averaged.

### Processing of single-cell RNA-seq datasets from GBM *in vivo*

Raw read count matrices and meta-data were downloaded from Neftel et al ^15^ and Muller et al ^54^ were downloaded from GSE131928 and UCSC Single Cell Browser (https://cells.ucsc.edu/), respectively. A Seurat object was created for each matrix separately and datasets were scaled. Subtype marker genes were downloaded from each dataset to maintain originally documented cell type annotations. In the Neftel dataset^15^, AHR-positive cells were denoted by cells with expression level > 2. Pair-wise matched expression of genes from each subtype were compared and statistical significance was determined by Wilcoxon-test. For the Muller dataset^54^, we annotated all cell types according to their highest expressed marker genes reported by the paper. We then used Seurat to generate violin plots for individual genes downregulated after AHR antagonism.

### Retrospective analysis of gene expression in human gliomas

Gene expression correlations of the AHR network were determined across primary patient gliomas and subtypes of GBM tumors, determined through analysis of the National Cancer Institute Repository for Molecular Brain Neoplasia Data (http://betastasis.com/glioma/rembrandt/) and TCGA (https://tcga-data.nci.nih.gov/publications/tcga), respectively. Individual gene expression survival analyses were performed for AHR network genes in primary patient GBM tumors using TCGA. High expression was defined as above the median and low expression below the median. Gene expression localization in structures of primary patient GBM was determined through analysis of Allen Institute of Ivy GAP (http://glioblastoma.alleninstitute.org/). Expression data was downloaded and heatmaps were generated using the matrix visualization software, Morpheus. Heatmaps of Pearson similarity matrices were also generated using Morpheus. Detailed information of heatmaps, including gene names in retained order as in **Figs. 5 and Supplementary 12b, 12d** can be found in **Supplementary Tables 2, 3 and 4.**

### Metabolic analysis of GBMO with Seahorse technology

For Seahorse Analysis (XFe96, Agilent Technologies), organoids were first dissociated via dissociation reagent. Dissociated cells were washed into warmed Seahorse XF DMEM medium supplemented with 10 mM glucose, 1 mM pyruvate, and 2 mM glutamine and plated at a density of 1x10^5^ cells/well on a poly-L-lysine coated XFe96 Seahorse cell culture microplate. Cells were simultaneously tested for oxygen consumption rate (OCR) and extracellular acidification rate (ECAR) per the manufacturer’s XF Real-Time ATP Rate Assay Kit protocol. Mitochondrial ATP (mitoATP) and glycolytic ATP (glycoATP) production rates were calculated via Agilent Seahorse XF Real-Time ATP Rate Assay Report Generator. ATP Production Rates were analyzed via t-test or ANOVA with Bonferroni posthoc corrections as needed with significance set at p<0.05.

### Organoid and GSCs treatment with AHR agonist and antagonist

To determine the effect of AHR treatment on GSC-derived organoid size, organoids were cultured as previously shown. After 4 days of stationary growth, the tissue droplets were transferred to a spinning bioreactor containing organoid differentiation media including 48% Neurobasal, 48% DMEM/F12, 1% B27 supplement, 0.5% N2 supplement, 1% Glutamax, 0.5% MEM-NEAA, and 1% Antibiotic-Antimycotic^45^ then cultured in the presence of the AHR agonist 6-formylindolo-3,2-b-carbazole (FICZ, Abcam), AHR antagonist 6, 2’, 4’-trimethoxyflavone (TMF, MilliporeSigma), and AHR antagonist CH223191 (Millipore Sigma) at each concentration for a week. The sizes of the organoids were determined under inverse microscope (U-TBI90, Olympus, Tokyo, Japan). Each experiment was performed three times with comparable results. GBMO organoids were measured after acquisition of images.

### Limiting dilution, self-renewal analysis after AHR treatment of GSCs

Cells were grown in 96-well plates. For the limiting dilution, cells were plated and wells were evaluated for number of tumorspheres at least 25 µm in size (sphere forming frequency). The percent of empty wells (with less than 75% growth number of wells with tumorspheres less than the total number of cells plated) were enumerated and the negative log (-log) of the proportion of negative wells was calculated for each cell density. This result was plotted to determine the clonal frequency of the culture conditions.

### CRISPR-Cas9 AHR inactivation of GSCs and GBMOs

To inactivate AHR in GBM-30, lentivirus for AHR sgRNA CRISPR (ABM, Canada) were infected into GBM-30 cells according to the manufacturer’s protocol. For viral infection, cells were plated and cultured overnight, and viral supernatants were added with polybrene (Millipore, Billerica, MA, USA) at a final concentration of 1 mg/mL. Cells were infected for at least 6 hours and allowed to recover for 24 hours with fresh medium. Infected cells were selected with puromycin 1 μg/mL for 48 hr. To confirm genetic engineering, cells were prepared for sequencing. 100% of the clones showed mutation and 80% were frameshift in exon 1 of the AHR gene.

### Individual gene Pearson’s correlation with AHR

To test the pair-wise correlation of AHR expression with other genes in GBM, all available RNA- seq data of primary patient GBM tumors were downloaded from TCGA. The expression data were mined for AHR-network genes and the top-most dysregulated genes detected by microarray. Pair-wise Pearson correlations and linear regressions were performed using GraphPad Prism (GraphPad Software).

### Survival analysis based on AHR expression signature in GBM

The identification of AHR-related gene signatures for survival prediction was performed on expression data acquired from the TCGA repository. From the GBM data set, we selected a subset of the most downregulated genes detected in the AHR antagonist-treated GBM organoids and randomly assigned the samples to a training set (350 subjects) and a testing set (174 subjects). Genes in the training set were prioritized using Cox regression with three models: LASSO, ElasticNet, and Ridge regression, as described in^55^. The results were then validated in the testing set and the prediction performance was assessed using Uno’s estimator of cumulative/dynamic AUC for right-censored time-to- event data (Uno’s AUC) and *c*-index^56^ For the best performing model, we identified sub-groups of subjects with favorable and poor prognosis defined as high- and low-risk groups, then plotted the Kaplan- Meier curves and performed the Mantel-Cox log-rank test. The analyses were performed using R programming language.

### Statistical analysis

All data were included for statistical analyses using GraphPad Prism 6.0. Unpaired Student’s t- test (two-tailed) was used for the comparison between two unpaired groups and one-way ANOVA was applied for multi-group data comparisons. The variance was similar between the groups that were being statistically compared. All data met the assumptions of the tests. Survival estimates were calculated using the Kaplan–Meier analysis. Briefly, the expression levels of target genes and patient survival information from the TCGA database were loaded into X-Tile as a tab-delimited text file. By running ‘Kaplan–Meier’ program, the cohort was then divided into two data sets with the optimal cut points generated according to the highest w2-value defined by log-rank test and Kaplan–Meier analyses. Bar graphs were presented as means ± SEM. with statistical significance at *P < 0.05, **P < 0.01, or ***P < 0.01.

## EXTENDED DATA

**Supplementary Figure 1.**
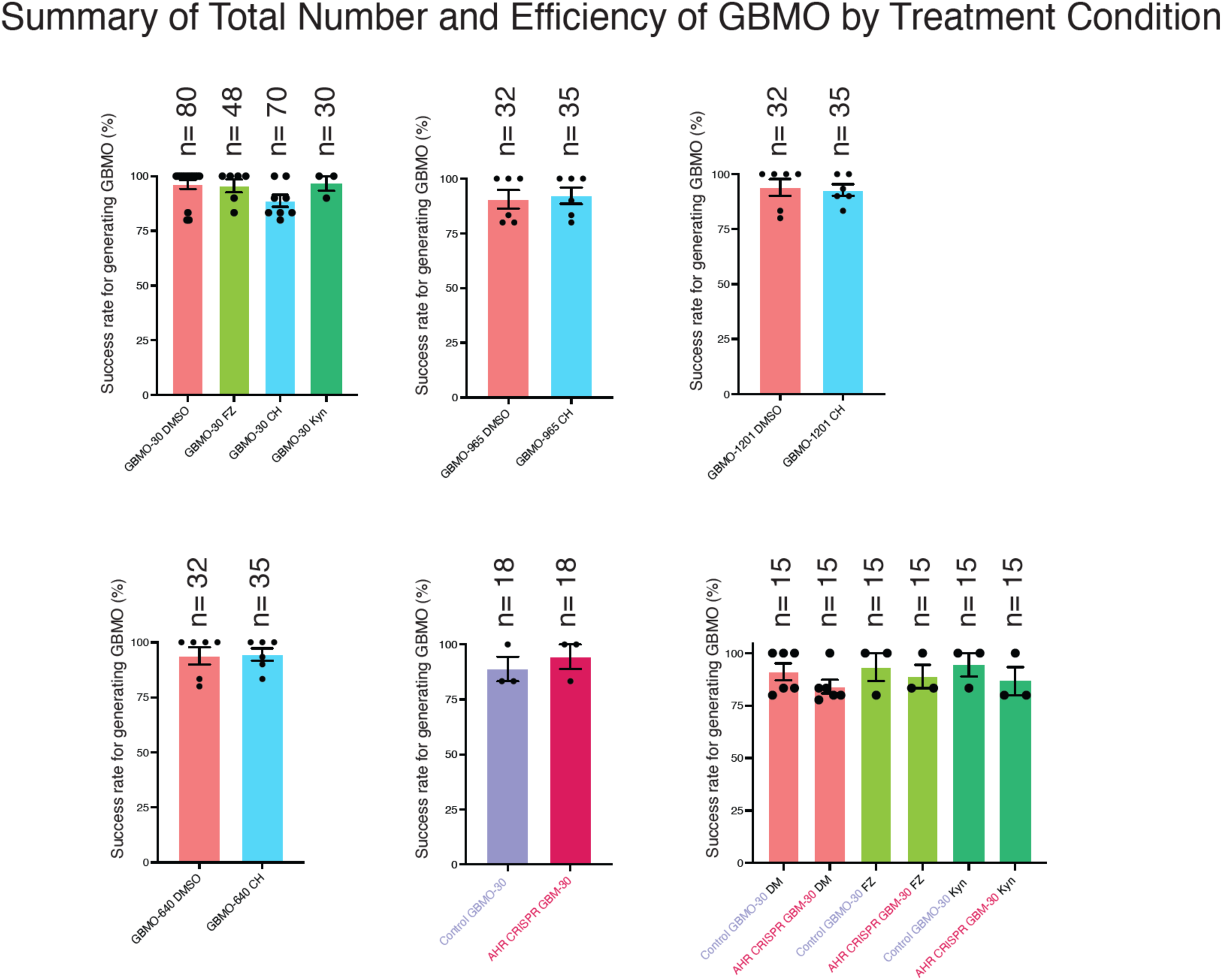
Summary figure of number of organoids efficacy. A total of 550 organoids were used as in methods with an efficiency of 95%.

**Supplementary Figure 2.**
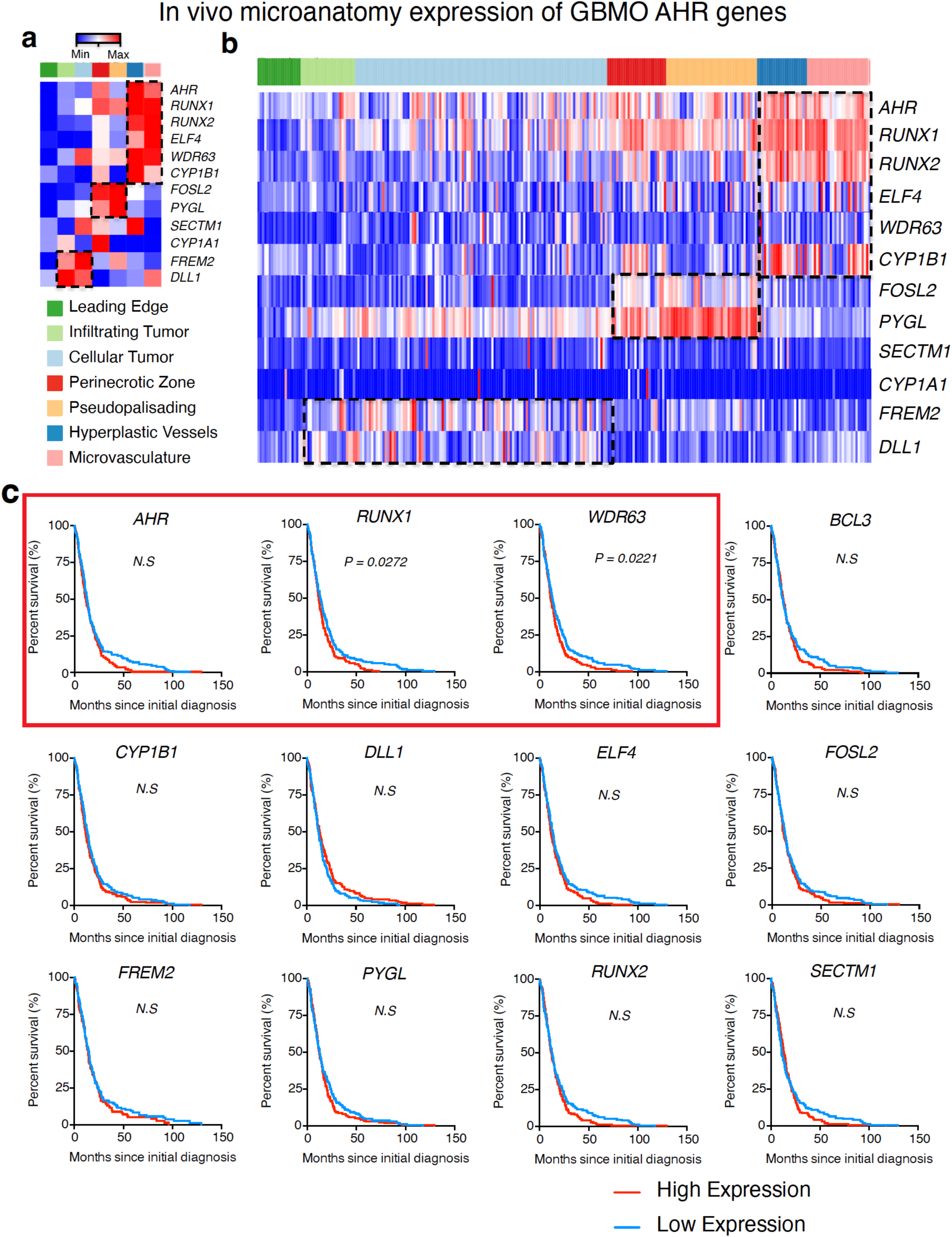
In vivo GBM microstructure localization, population survival curves, (Related to Figure 1). **(a)** Heatmap showing average expression of AHR-network genes by tumor structure. **(b)** Complete heatmap depicting expression of AHR-network genes in tumor structures. Tumor samples comprise the x-axis. RNA-Seq data of tumor structures are from IvyGAP. **(c)** Kaplan-Meier curves of patients with GBM distinguished by the expression of AHR-network genes. Red: high expression; blue: low expression. Survival and RNA-Seq data derived from publicly available TCGA cohorts; Mantel-Cox analysis. (*RUNX1*: *P* = 0.0272; *WDR63*: *P* = 0.0221.)

**Supplementary Figure 3.**
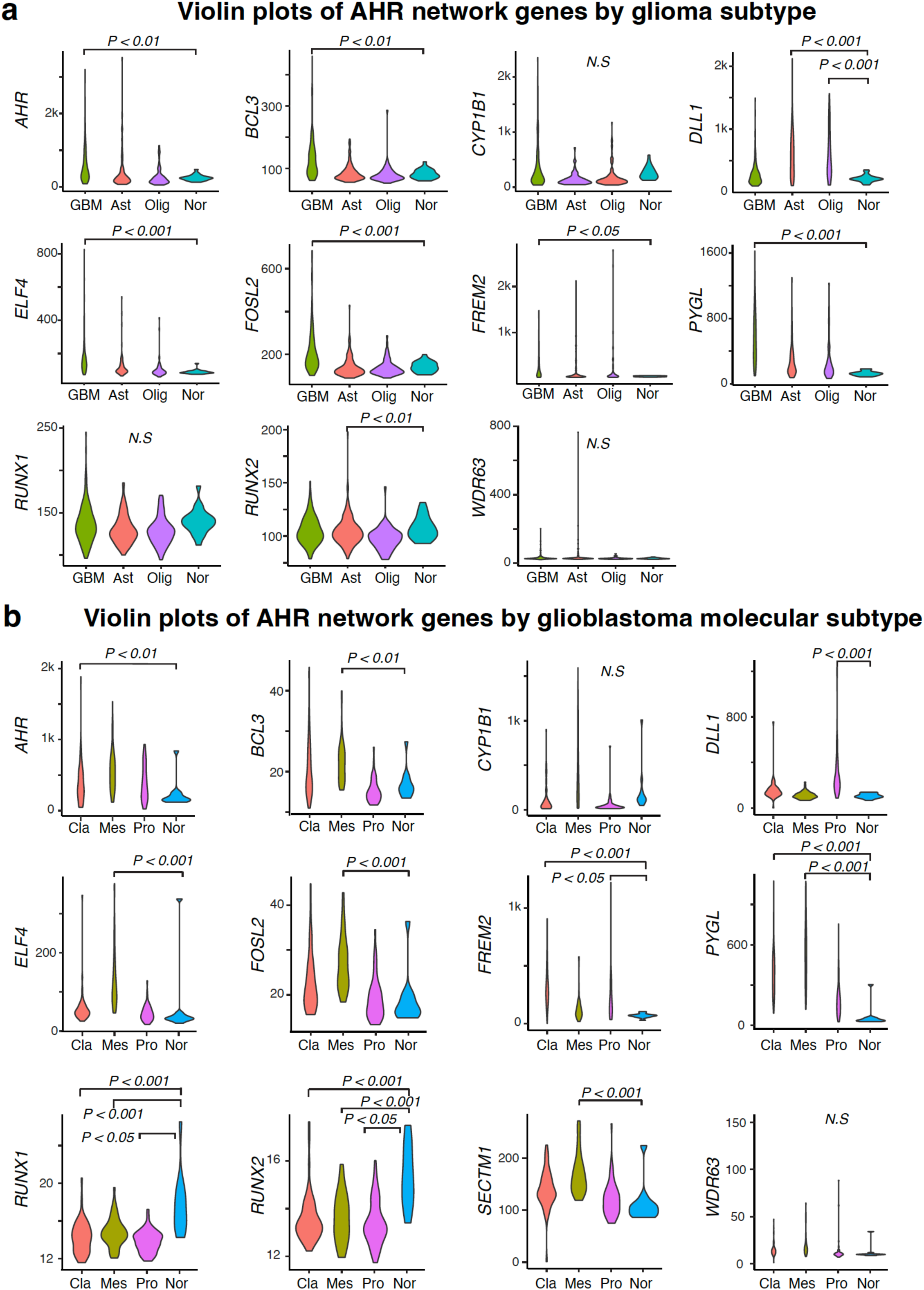
in vivo expression of AHR network genes in gliomas and GBM subtypes. (Related to Figure 1). **(a)** Violin plots of AHR network gene expression by glioma type, using publicly available TCGA cohorts. GBM, Glioblastoma; Ast, Astrocytoma; Olig, Oligodendroglioma; Nor, Normal. One-way ANOVA followed by Dunnett’s correction, relative to Normal. (*AHR*: GBM, *P* < 0.01. *BCL3*: GBM, *P* < 0.01. *DLL1*: Ast, *P* < 0.001; Olig, *P* < 0.001. *ELF4*: GBM, *P* < 0.001. *FOSL2*: GBM, *P* < 0.001. *FREM2*: GBM, *P* < 0.05. *PYGL*: GBM, *P* < 0.001. *RUNX2*: Ast, *P* < 0.01.) **(b)** Violin plots of AHR network gene expression by GBM subtype, using publicly available TCGA cohorts. Cla, Classical; Mes, Mesenchymal; Pro, Proneural; Nor, Normal. One-way ANOVA followed by Dunnett’s correction, relative to Normal. (*AHR*: Cla, *P* < 0.01. *BCL3*: Mes, *P* < 0.01. *DLL1*: Pro, *P* < 0.001. *ELF4*: Mes, *P* < 0.001. *FOSL2*: Mes, *P* < 0.001. *FREM2*: Cla, *P* < 0.001; Pro, *P* < 0.05. *PYGL*: Cla, *P* < 0.001; Mes, *P* < 0.001. *RUNX1*: Cla, *P* < 0.001; Mes, *P* < 0.001; Pro, *P* < 0.05. *RUNX2*: Cla, *P* < 0.001; Mes, *P* < 0.001; Pro, *P* < 0.05. *SECTM1*: Mes, *P* < 0.001). N.S = not significant, *P* > 0.05.

**Supplementary Figure 4.**
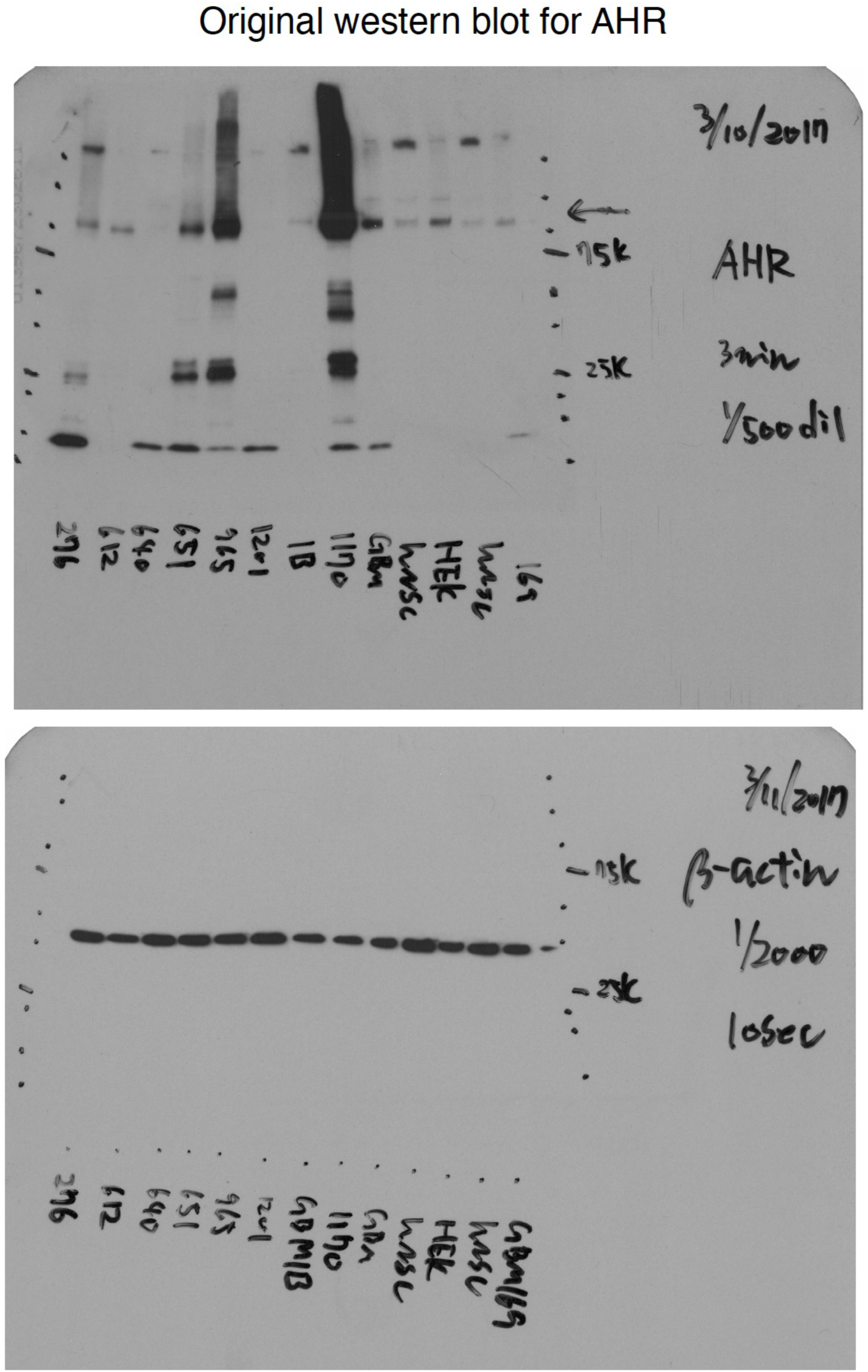
Full-Sized Western Blots. Corresponding to AHR (upper) and β-actin (lower) blots from Figure 3k with Ladder Indicating Molecular Weight Values **(<u>Related to</u>** Figure 1).

**Supplementary Figure 5.**
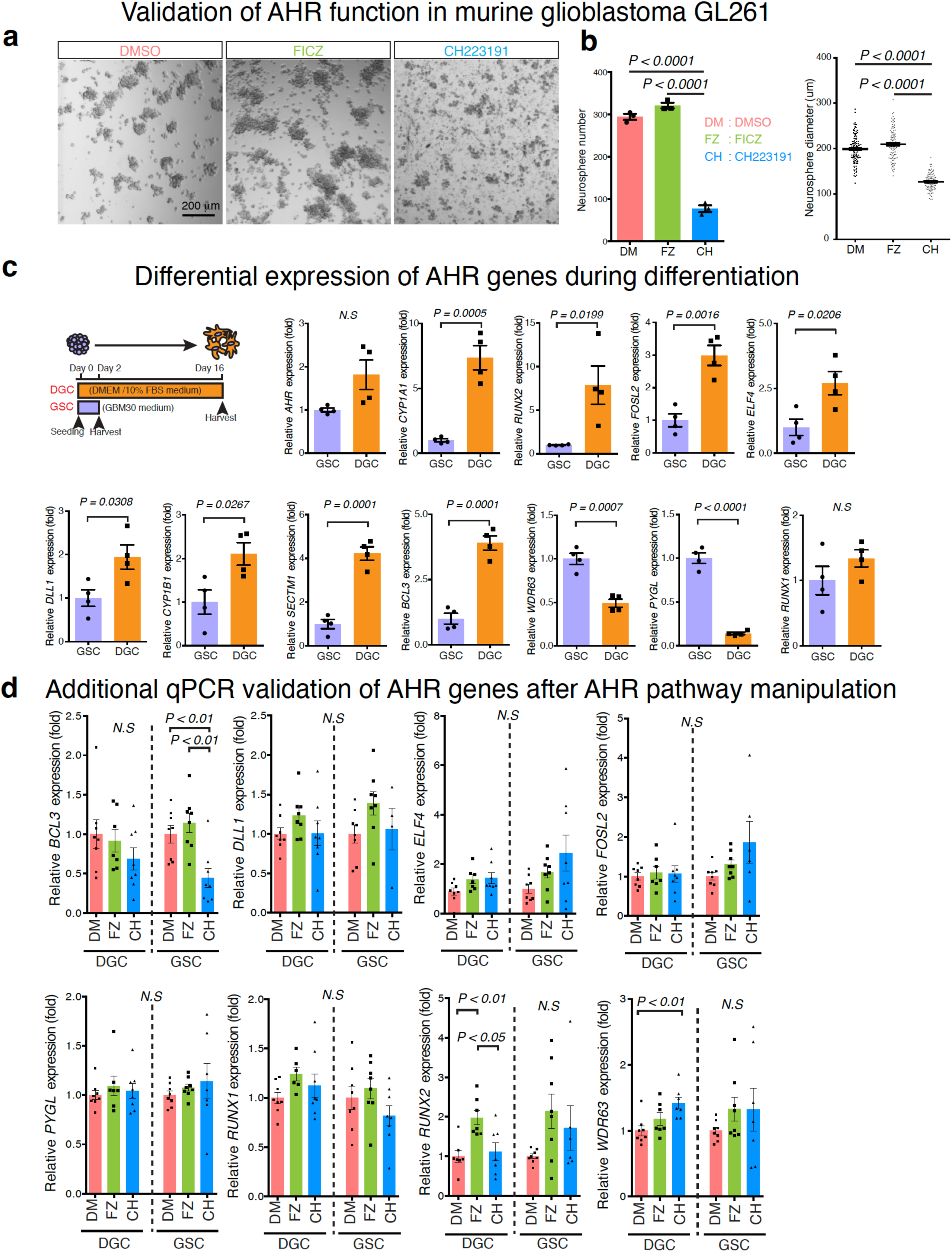
Mouse GSC self-renewal with AHR targeting and AHR network gene expression across GSC differentiation (Related to Figure 2). **(a)** Sample bright-field images of mouse glioma GL261N tumorspheres after treatment with DMSO, FICZ 500 nM), and CH223191, 3 mM) for 7 days. Scale bar, 200 μm. **(b)** Quantitative analysis of neurosphere number and diameter after treatment with AHR exogenous ligands. One-way ANOVA with Tukey’s test; *n* = 3 independent experiments. Data are presented as mean ± SEM. (Number: DMSO vs. FICZ, *P* < 0.0001; FICZ vs. CH223191, *P* < 0.0001. Diameter: DMSO vs. FICZ, *P* < 0.0001; FICZ vs. CH223191, *P* < 0.0001). **(c)** Time course and culture conditions for differentiated glioblastoma cells (DGCs) and non-differentiated GSCs. Quantitative RT-PCR for AHR effectors in GSCs and DGCs under normal conditions. Two-tailed unpaired Student’s t- tests; *n* = 3 independent experiments. Data are presented as means ± SEM. (*CYP1A1*, *P* = 0.0005; *CYP1B1*: *P* = 0.0267; *SECTM1*, *P* = 0.0001; *BCL3*, *P* = 0.0001; *RUNX2*, *P* = 0.0199; *FOSL2*, *P* = 0.0016; *ELF4*, *P* = 0.0206; *DLL1*, *P* = 0.0308; *WDR63*, *P* = 0.0007; *PYGL*, *P* < 0.0001). **(d)** Differential targeting of AHR in differentiated glioblastoma cells (DGCs) or non-differentiated GSCs. One-way ANOVA with Tukey’s test; *n* = 3 independent experiments. Data are presented as mean ± SEM. (*BCL3* GSC: DMSO vs. CH223191, *P* < 0.01; FICZ vs. CH223191, *P* < 0.01. *RUNX2* DGC: DMSO vs. FICZ, *P* < 0.01; FICZ vs. CH223191, *P* < 0.05. *WDR63* DGC: DMSO vs. CH223191, *P* < 0.01.). N.S. = not significant, *P* > 0.05.

**Supplementary Figure 6.**
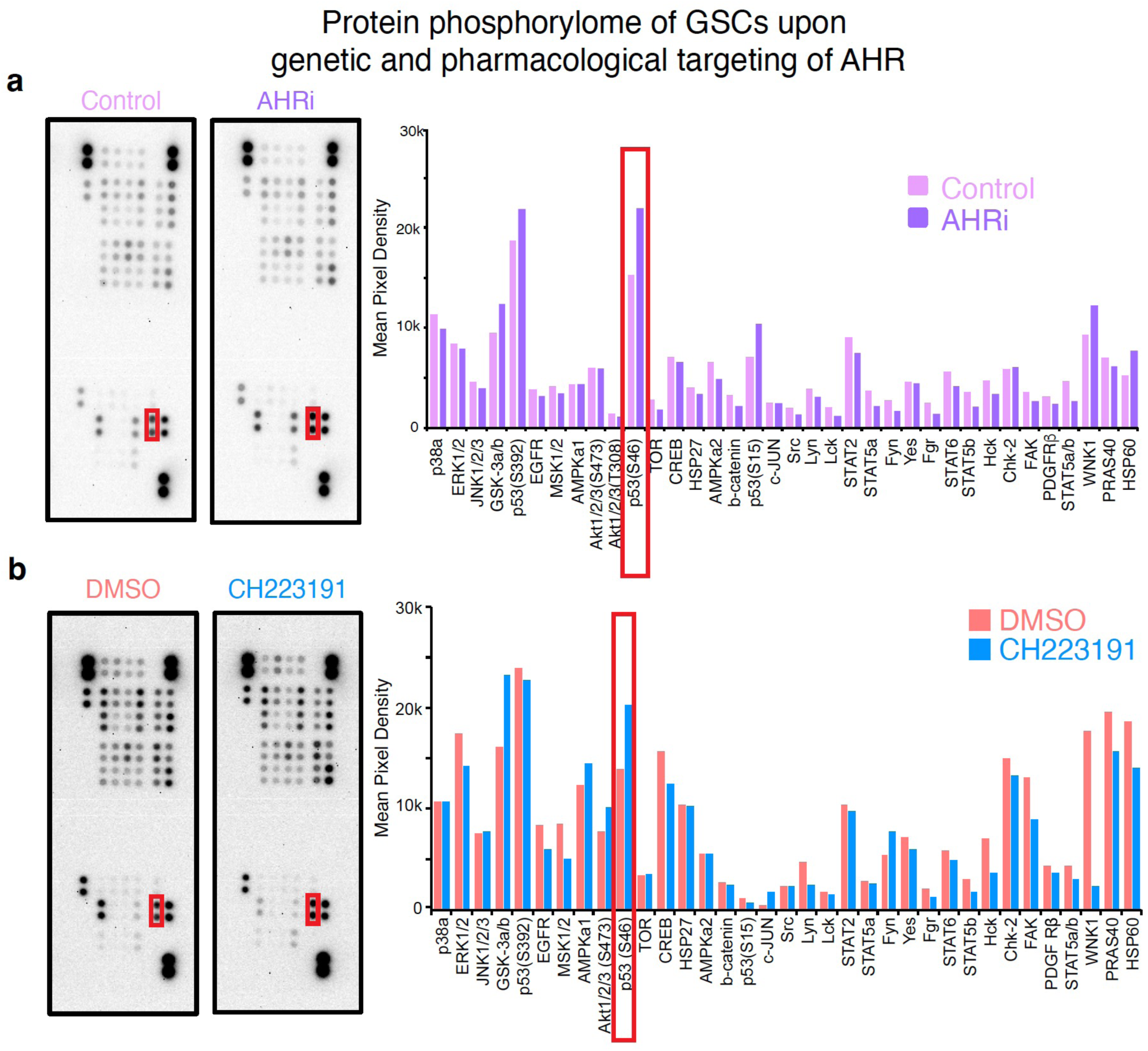
Protein phosphorylome of GSCs upon genetic and pharmacological targeting of AHR (Related to Figures 2). **(a)** Image of antibody array and mean pixel density for GSCs transfected with mock siRNA (Control) or siRNA targeting AHR transcripts (AHRi). Red boxes indicate the expression of p53 (S46). **(b)** Image of antibody array and mean pixel density for GSCs treated with DMSO and AHR antagonist, CH223191. Red boxes indicate the expression of p53 (S46).

**Supplementary Figure 7.**
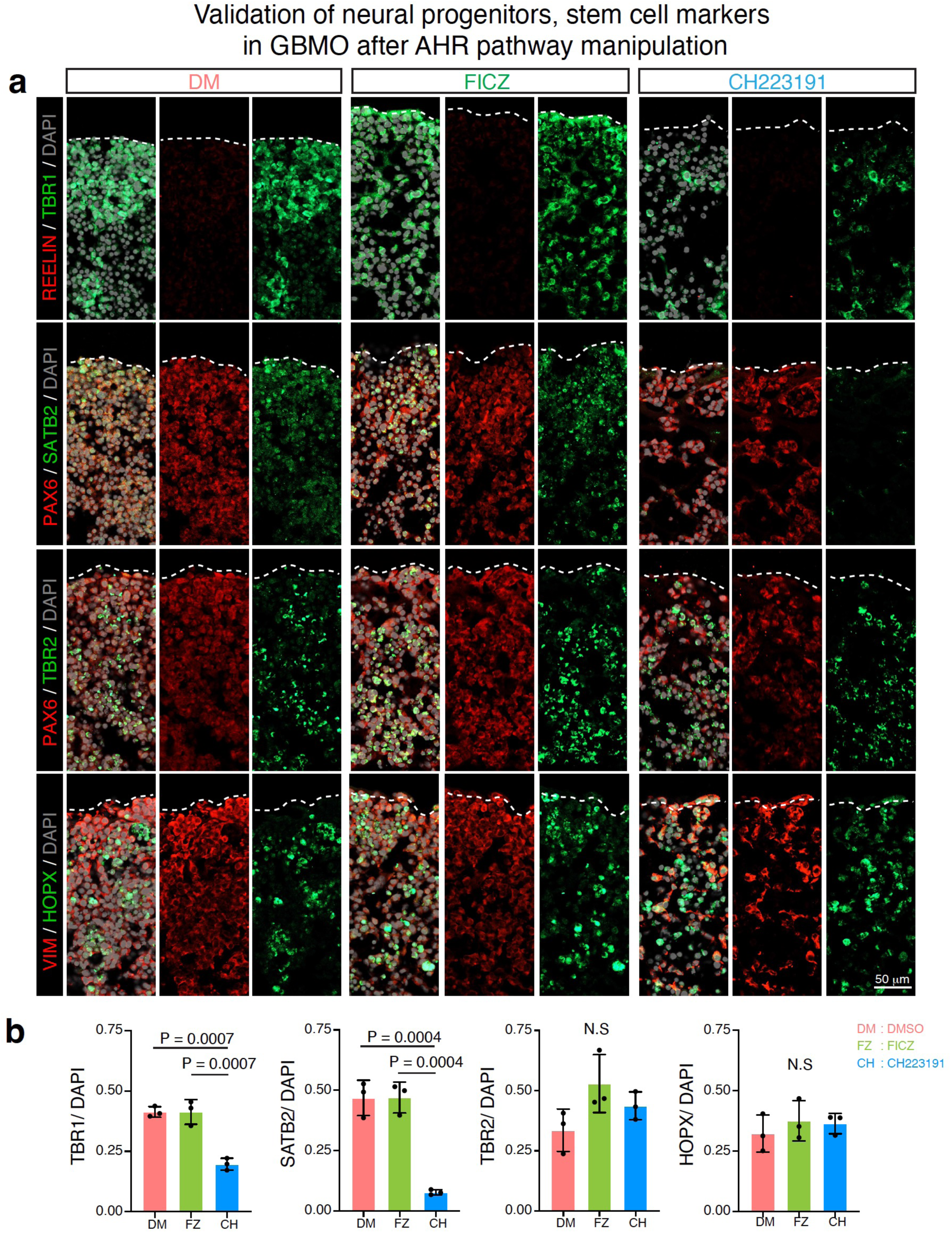
Progenitor marker expression in AHR drug-treated GBMO-30 (Related to Figures 3). **(a)** Representative confocal images for immunostaining of Cajal-Retzius cell (top: REELIN), deep-layer cortical neuron (top: TBR1, second: SATB2), intermediate progenitor (second: TBR2), radial glia (second and third: PAX6, bottom: VIM), and outer radial glia (bottom: HOPX) markers in GBMO-30 after DMSO, FICZ, and CH223191 treatment. Scale bar, 50 μm. **(b)** Quantification of TBR1 (top), SATB2 (second), TBR2 (third), and HOPX (bottom)-positive cells in GBMO-30 after DMSO, FICZ, and CH223191 treatment. Bar graphs are presented as means ± SEM. Two-tailed unpaired *Student’s t-test*. (TBR1: DMSO vs. FICZ, *P* = 0.0007; FICZ vs. CH223191, *P* < 0.0007. SATB2: DMSO vs. FICZ, *P* = 0.0004; FICZ vs. CH223191, *P* < 0.0004.). N.S. = not significant, *P* > 0.05.

**Supplementary Figure 8.**
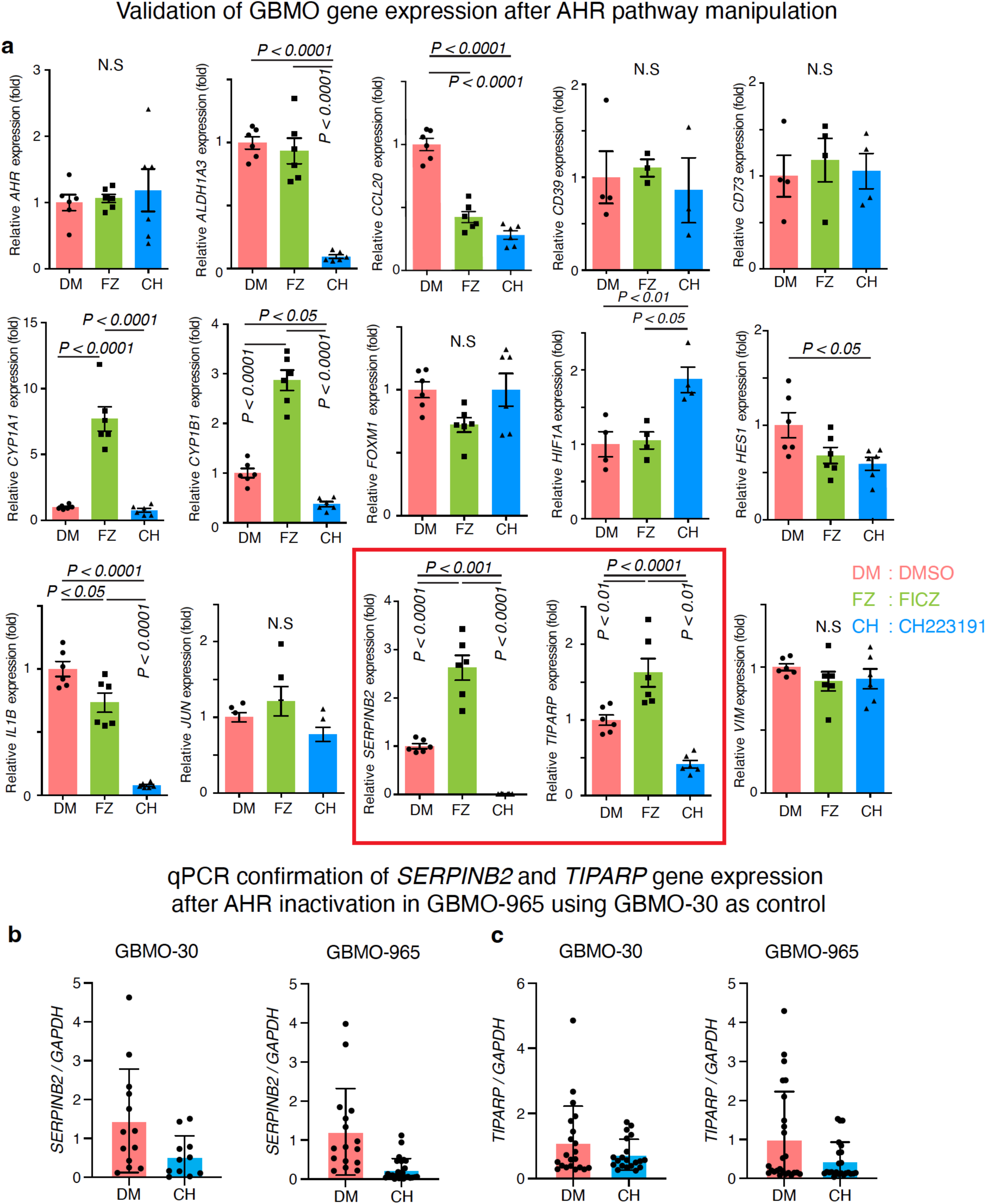
Extended analysis of AHR drug-treated GBMO-30 transcriptomes and confirmatory qPCR expression (Related to Figure 5). **(a)** Confirmatory qPCR for genes related to stemness and immunity in AHR drug-treated GBMO-30. One-way ANOVA with Tukey’s test; *n* = 3 independent experiments. Data are presented as mean ± SEM. (*CYP1A1*: DMSO vs. FICZ, *P* < 0.0001; FICZ vs. CH223191, *P* < 0.0001. *CYP1B1*: DMSO vs. FICZ, *P* < 0.0001; DMSO vs. CH223191, *P* < 0.05; FICZ vs. CH223191, *P* < 0.0001. *ALDH1A3*: DMSO vs. FICZ, *P* < 0.0001; FICZ vs. CH223191, *P* < 0.0001. *CCL20*: DMSO vs. FICZ, *P* < 0.0001; FICZ vs. CH223191, *P* < 0.0001. *HES1*: DMSO vs. CH223191, *P* < 0.05. *IL1B*: DMSO vs. FICZ, *P* < 0.05; DMSO vs. CH223191, *P* < 0.0001; FICZ vs. CH223191, *P* < 0.0001. *SERPINB2*: DMS0 vs. FICZ, *P* < 0.0001; DM vs. CH223191, *P* < 0.0001; FICZ vs. CH223191, *P* < 0.0001. *TIPARP*: DMS0 vs. FICZ, *P* < 0.0001; DMSO vs. CH223191, *P* < 0.0001; FICZ vs. CH223191, *P* < 0.0001. *HIF-1*α: DMSO vs. CH223191, *P* < 0.01; FICZ vs. CH223191, *P* < 0.05). **(b, c)** Expression of selected AHR network genes (SERPINB2: b and TIPARP in c in GBMO-965 (right) using as control GBMO-30 (left) during inactivation with CH223191.

**Supplementary Figure 9.**
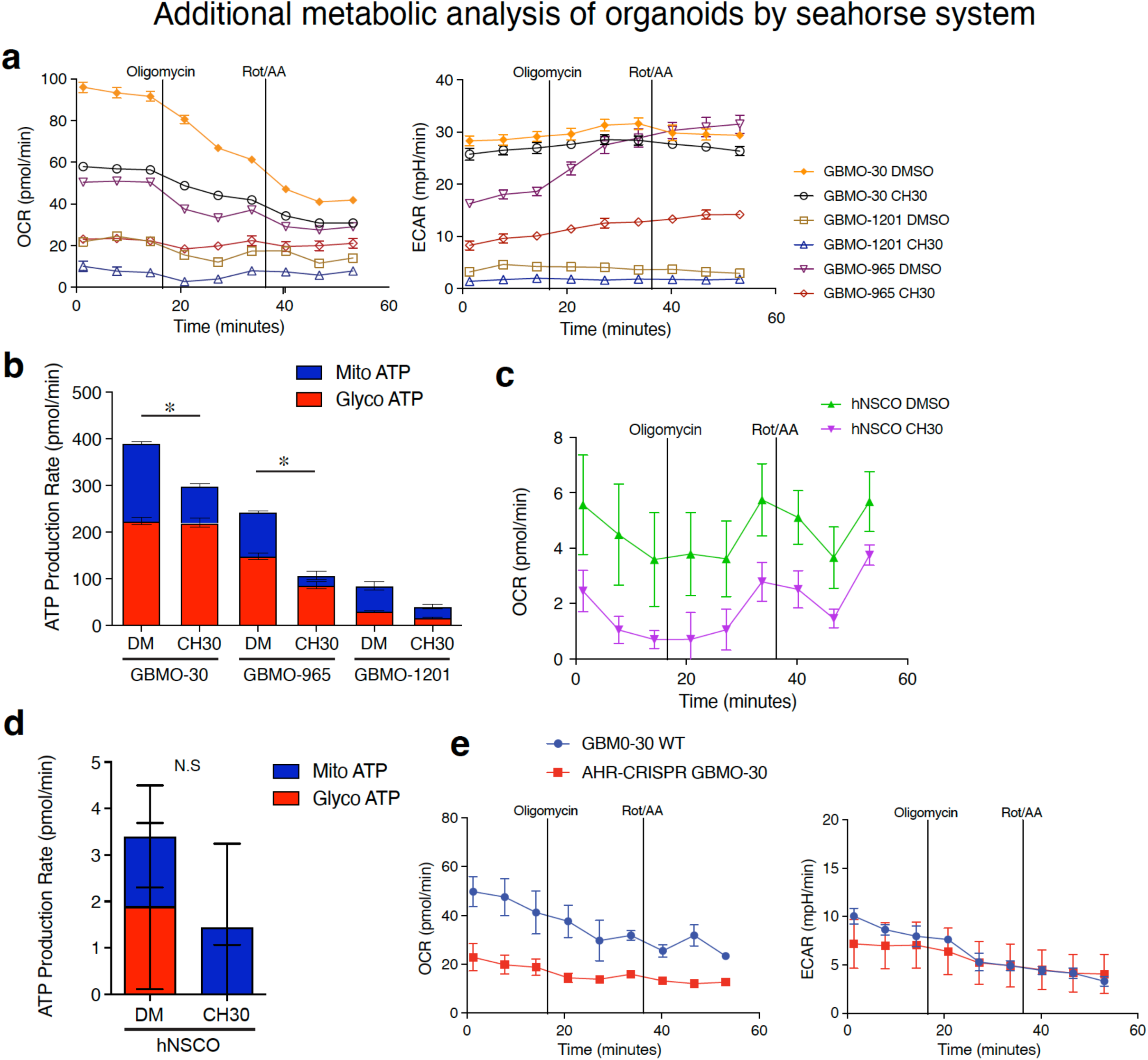
Seahorse XF Real-Time ATP Rate Assay for AHR-CRISPR GBMO-30 and CH223191 treated GBMOs (Related to Figure 5). **(a)** Measurement for Oxygen Consumption Rate (OCR, left) and Extracellular Acidification Rate (ECAR, right) in GBMO-30, -965, and -1201 with 30 μM CH223191. **(b)** Mitochondrial ATP (Mito ATP) and glycolytic ATP (Glyco ATP) production rates in each CH223191-treated GBMO. **(c)** OCR in CH223191-treated hNSCO. **(d)** Mito ATP and glyco ATP production rates in CH223191-treated hNSCO. **(e)** OCR (left) and ECAR (right) in AHR-CRISPR GBMO- 30.

**Supplementary Figure 10.**
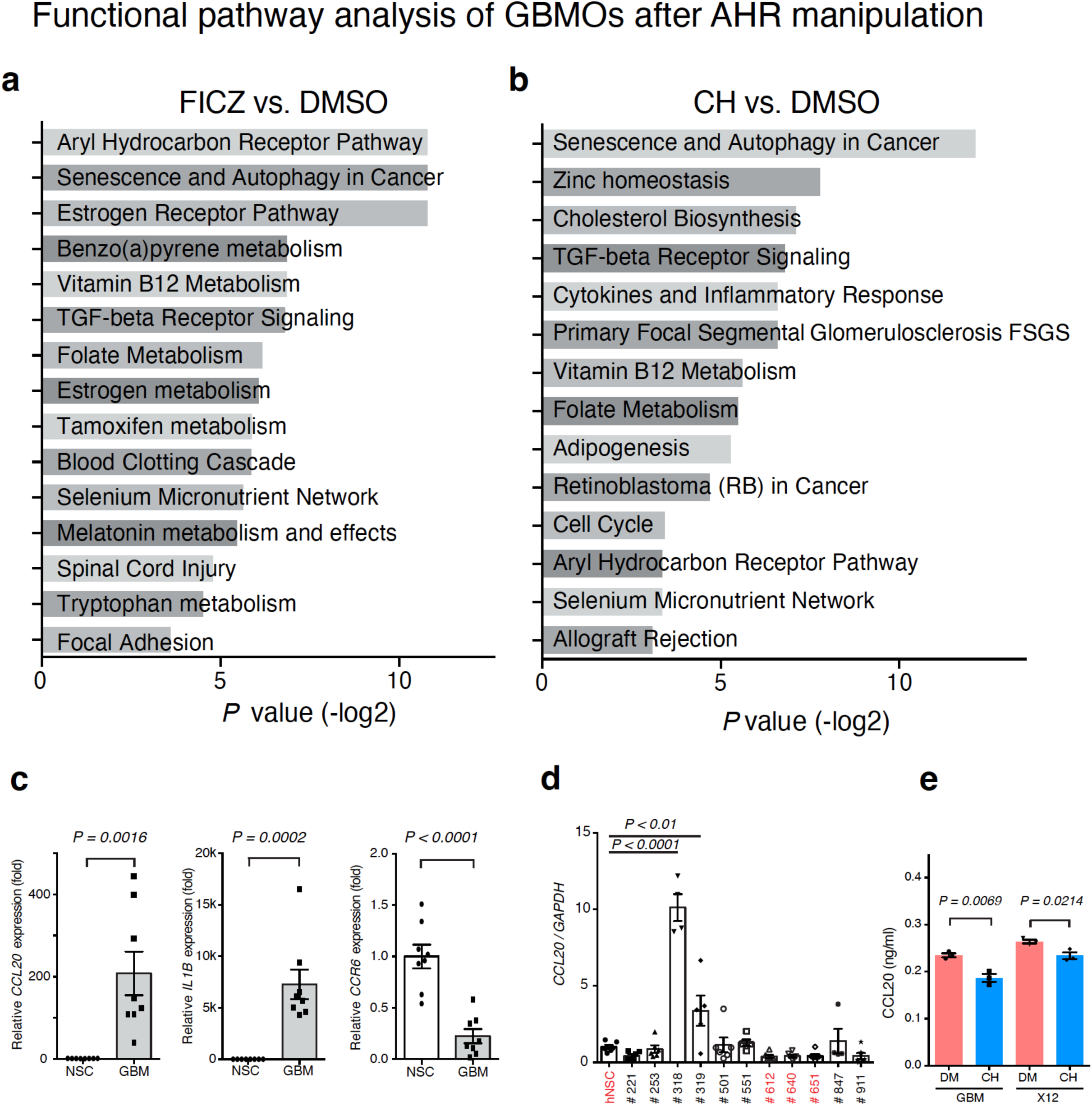
Pathway analysis of DEGs and analysis of AHR target CCL20 and IL1B in GBMO-30 (Related to Figure 5). **(a)** Pathway analysis of DEGs induced by treatment of GBMO-30 with FICZ indicates enrichment for metabolism and immune cell migration and adhesion. **(b)** Pathway analysis of DEGs induced by treatment of GBMO-30 with CH223191 reveals enrichment for immune signaling, apoptosis, and cell cycle. **(c)** mRNA expression of *CCL20*, *IL1B*, and *CCR6* in GSCs (GBM, grey bars), relative to NSCs (white bars). Two-tailed unpaired Student’s t-test; *n* = 3 independent experiments. Data are presented as means ± SEM. (*CCL20*, *P* = 0.0016; *IL1B*, *P* = 0.0002; *CCR6*, *P* < 0.0001.) **(d)** Quantitative RT-PCR for *CCL20* using multiple GBM cell lines with Dunnett’s correction, relative to hNSC; *n* = 3 independent experiments. Data are presented as means ± SEM. (GBM318, *P* < 0.0001; GBM319, *P* < 0.01.) **(e)** ELISA analysis for CCL20 of GBM30 (GBM) and X12 GSCs after treatment with DMSO and with CH223191.Two-tailed unpaired Student’s t-test; n = 3 independent experiments. Data are presented as means ± SEM. (GBM, *P* = 0.0069; X12, *P* = 0.0214.)

**Supplementary Figure 11.**
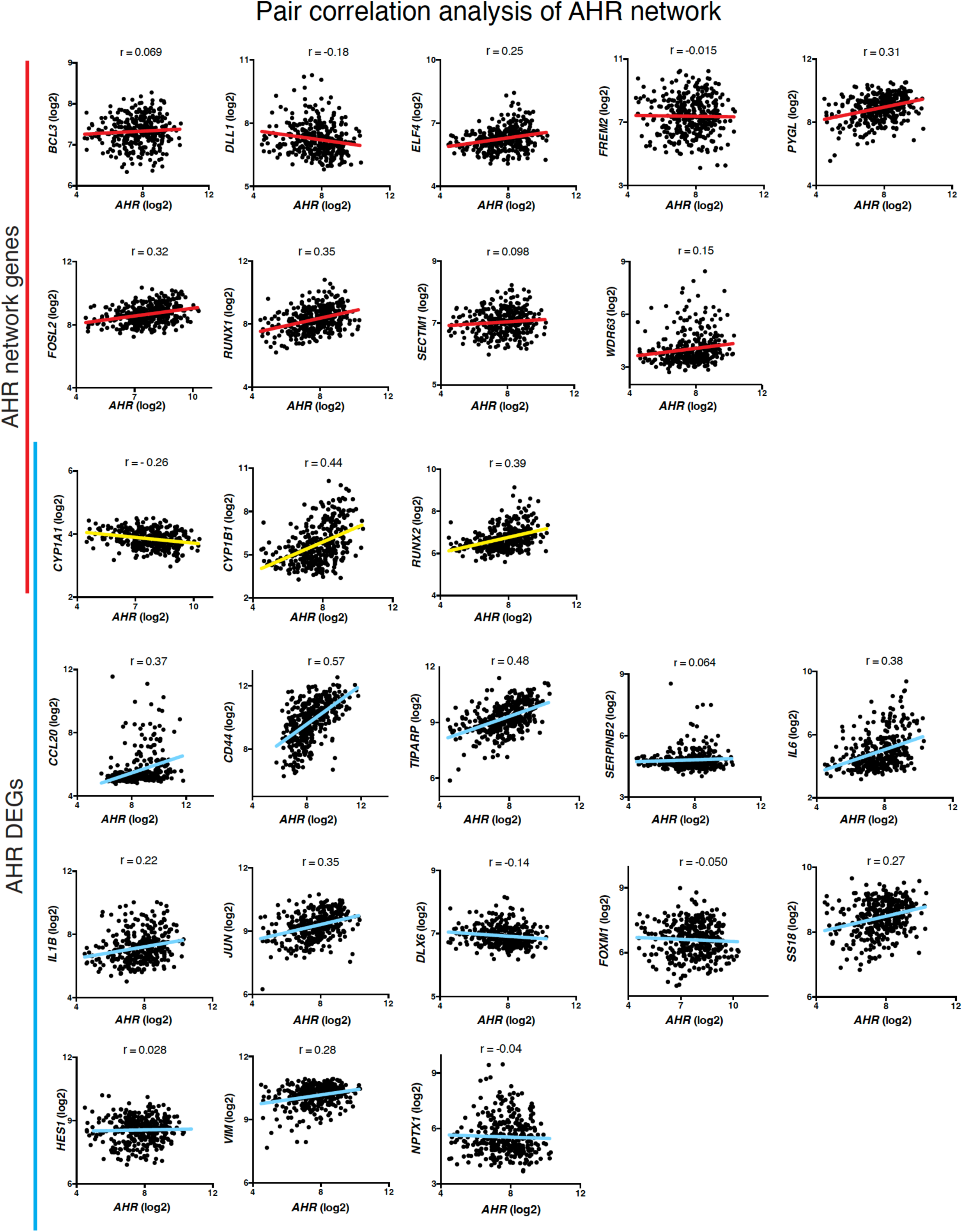
Pairwise correlation of AHR and downstream gene expression in GBM in vivo (Related to Figure 5). Pairwise correlation analyses between AHR, AHR-network genes (red), and DEGs identified by transcriptome analysis of with CH223191.-treated GBMO-30 (blue) in primary GBM samples from TCGA. Numbers are Pearson coefficient values scaled from 1 to -1 representing a positive and negative linear correlation, respectively, between the two genes expressed in GBM *in vivo*.

**Supplementary Figure 12.**
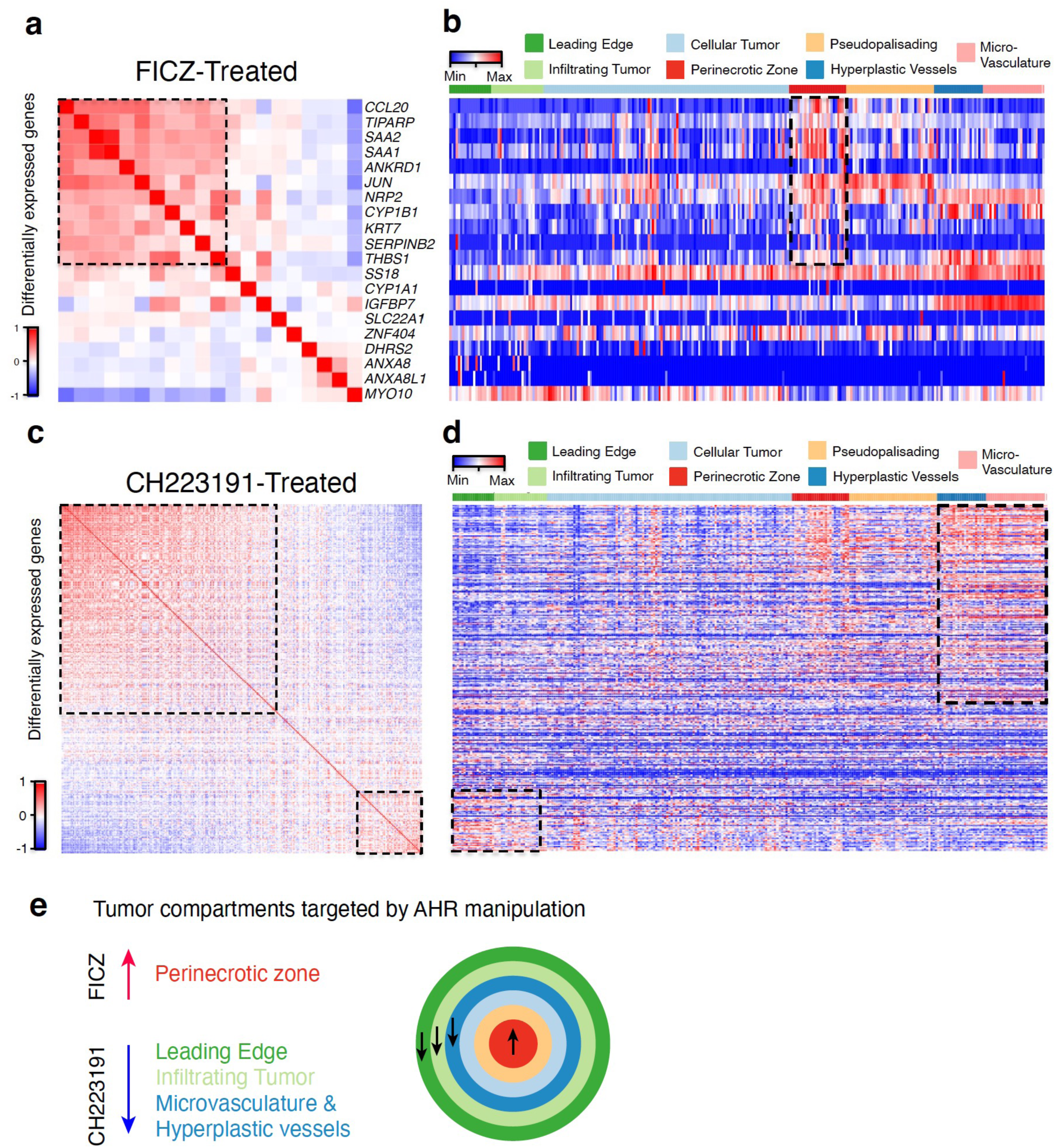
(Related to Figure 5). (a-d) Heatmaps depict similarity matrices (a, c) and GBM microstructure localization of DEGs (b, d) in FICZ- (a, b) and CH223191-treated (c, d) GBMO-30. (**e**) Summary of gene expression changes by tumor compartment.

**Supplementary Figure 13.**
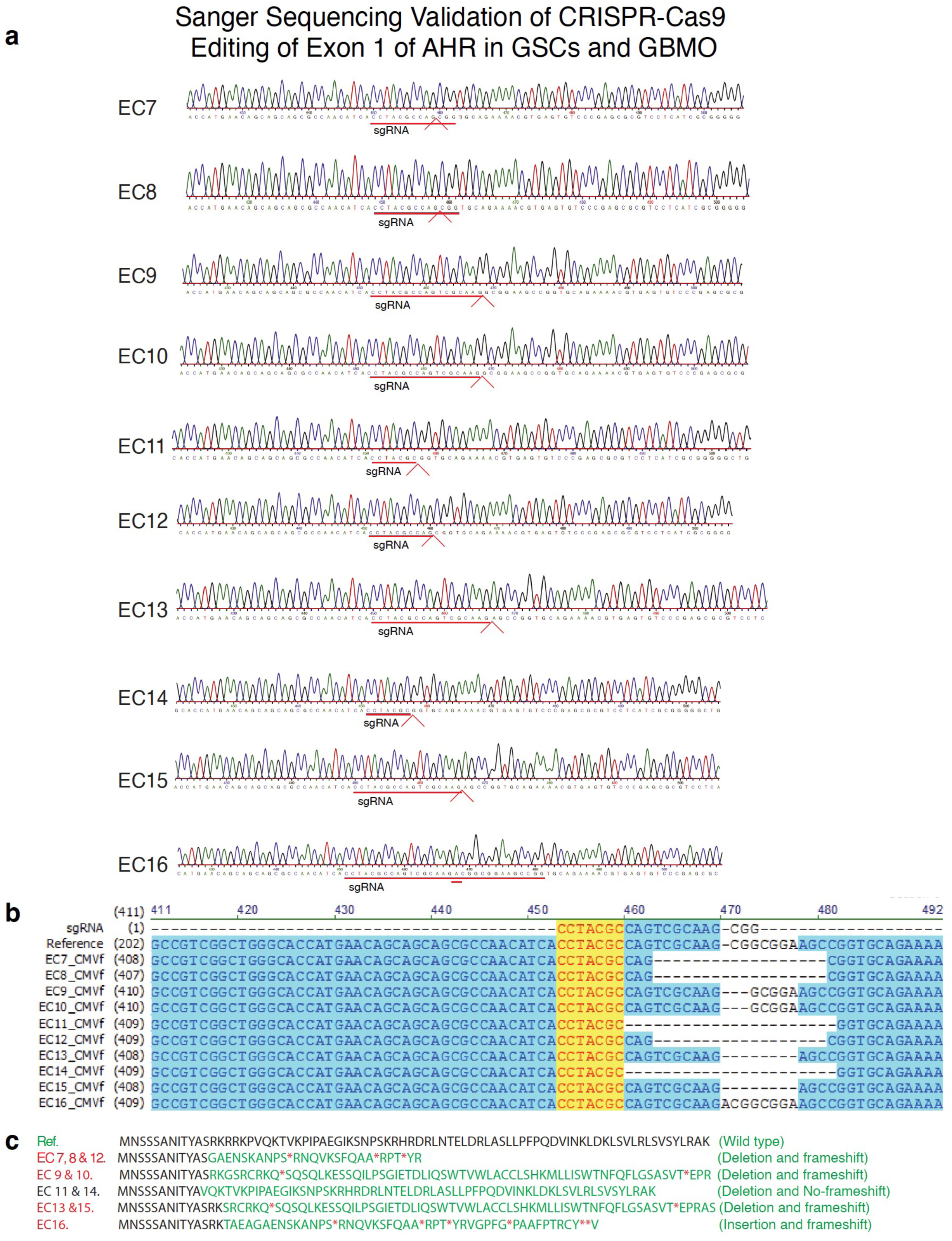
Sanger sequencing and validation of mutation for CRISPR edited GSCs and GBMOs (Related to Figure 6). **(a)** Subcloned DNA sequencings for target region in AHR exon 1. **(b)** Alignment of subcloned DNA sequencing for target region. **(c)** Predicted translation of AHR protein sequence from subcloned sequencing.

**Supplementary Figure 14.**
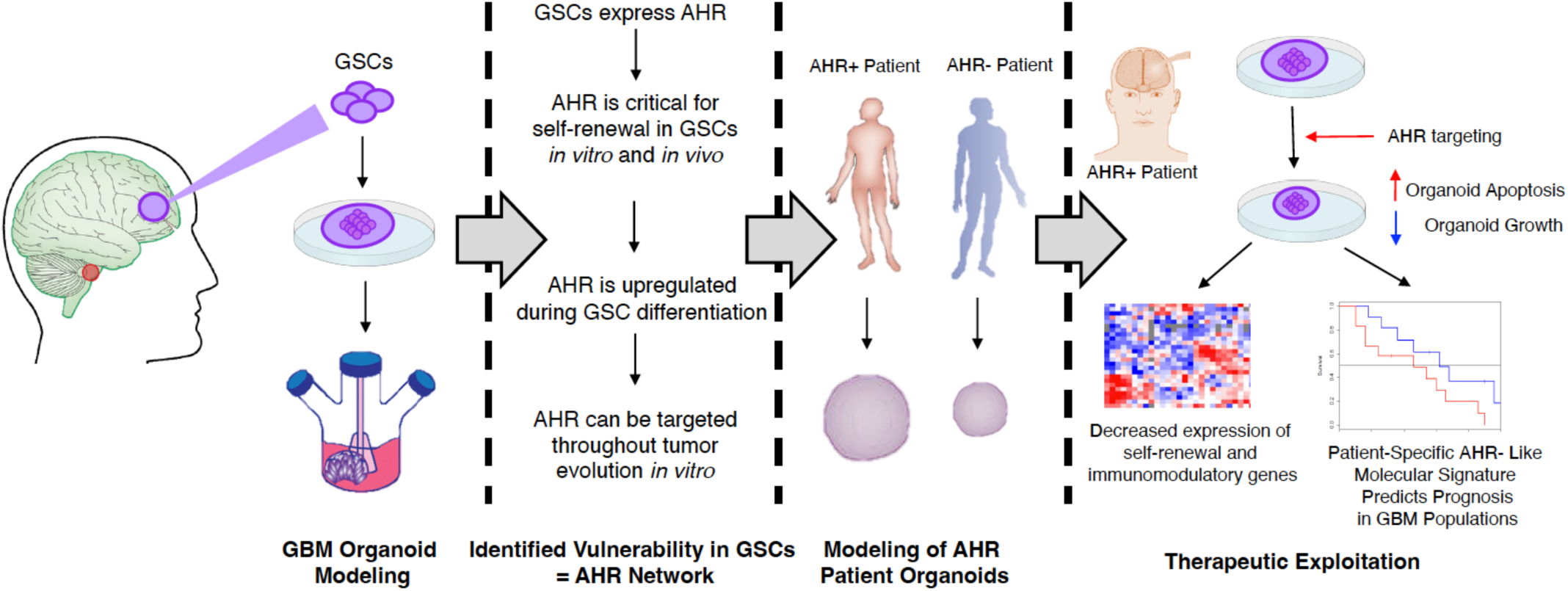
Summary schematic of our findings (Related to 7). **(a)** Patient-Derived GBMO Platform and Resulting Identification of AHR as a Molecular Vulnerability and Prognostically relevant Expression Signature.

**Supplementary Figure 15.**
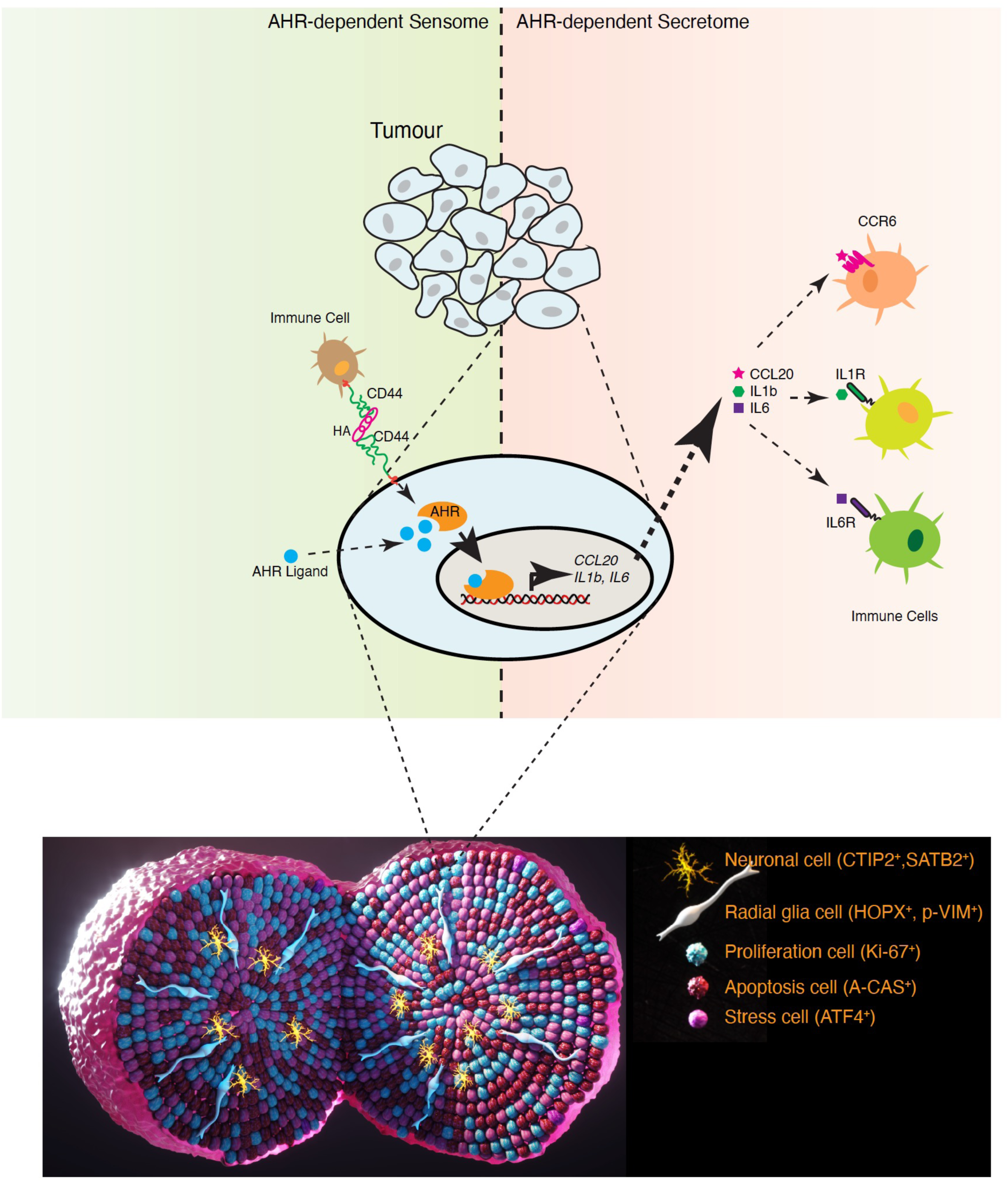
Model of mechanisms of Glioma-intrinsic AHR in GSCs and GBMO. Model of AHR-Mediated Effects on GBM Immunity within the TIME.

## Acknowledgments

We kindly thank Sarah Warner for technical expertise in microarrays. We thank Dr. Amy Lovett-Racke and Matthew Gormley for technical help. To Drs. Timothy P. Cripe MD, PhD, and Sofia B. Lizarraga, PhD for their critical reading of the manuscript. The work was supported in part by the NCI (CA016058) and P30 CA016058. AQH is supported by the NIH (CA200399, CA183827, CA195503, CA216855, NS070024), as well as the Mayo Clinician Investigator Award and the State of Florida. VP is supported by grant K24CA 160777. JI was supported by the NRI Research Award and The Biogen-UConn Collaboration grant. Dr. Imitola is a consultant for Biogen and Novartis.

## Notes

### Competing Interest Statement

The authors have declared no competing interest.

